# Analysis of (p)ppGpp metabolism and signaling using a dynamic luminescent reporter

**DOI:** 10.1101/2025.04.16.649062

**Authors:** Molly Hydorn, Sathya Narayanan Nagarajan, Elizabeth Fones, Caroline S. Harwood, Jonathan Dworkin

## Abstract

As rapidly growing bacteria begin to exhaust nutrients, their growth rate slows, ultimately leading to the non-replicative state of quiescence. Adaptation to nutrient limitation requires widespread metabolic remodeling that is in part mediated by the phosphorylated nucleotides guanosine tetra- and penta-phosphate, collectively (p)ppGpp. We have developed a novel reporter of (p)ppGpp abundance in the Gram-positive bacterium *Bacillus subtilis* based on the recent identification of a riboswitch that binds (p)ppGpp and modulates transcription via regulation of a transcriptional terminator. Placement of an unstable reporter, firefly luciferase, downstream of the riboswitch allows for sensitive and dynamic assessment of (p)ppGpp. We first confirm that the reporter accurately reflects (p)ppGpp abundance in a variety of well-established conditions. We then proceed to use it to demonstrate the physiological importance of several mechanisms of regulation of (p)ppGpp metabolism previously observed only *in vitro* including allosteric interactions between (p)ppGpp synthesis enzymes and the hydrolytic activity of a (p)ppGpp synthase. (p)ppGpp signaling has been implicated in the regulation of gene expression, and we demonstrate a close temporal association between gene expression and (p)ppGpp abundance, indicating a rapid, and therefore likely direct mechanism of (p)ppGpp dependent gene activation. Thus, this reporter provides a new, comprehensive analysis of (p)ppGpp signaling *in vivo* and offers the potential ability to sensitively monitor the temporal dynamics of (p)ppGpp abundance under diverse environmental conditions.

**Author Summary:** Most bacteria adapt to stressful conditions such as nutrient limitation by synthesizing a signaling molecule, known as ppGpp, that consists of a hyper-phosphorylated GTP. Synthesis of ppGpp affects most aspects of cellular physiology including replication, transcription and translation. We present here a method that for the first time allows measurement of ppGpp abundance in living cells, greatly facilitating investigation into ppGpp metabolism.

## Introduction

The adaptive response of bacteria to environmental changes can be relatively specific, as in the case of two-component signaling that involves a single transcription factor regulating a limited number of genes. Alternatively, an adaptive response can be broad, such as the widespread physiological changes often referred to as the “stringent response” mediated by the nucleotides guanosine tetraphosphate (ppGpp) and pentaphosphate (pppGpp), collectively (p)ppGpp. These molecules reduce replication, transcription, translation and GTP synthesis by direct inhibitory interactions with essential enzymes of these processes (1, 2). They also stimulate the expression of genes encoding proteins mediating stress protection (e.g., *B. subtilis hpf* (3)), but the mechanistic basis for this stimulation is not well understood.

In both Gram-positive and Gram-negative bacteria, RelA/SpoT Homolog (RSH) enzymes (4) synthesize (p)ppGpp in response to diminished amino acid availability as manifested by the presence of uncharged tRNA molecules in ribosome A-site (5). RSH enzymes contain a synthase domain that transfers the β- and γ-phosphate groups from ATP and adds them to the 3’ carbon of the ribose of a GTP or GDP molecule to form pppGpp or ppGpp, respectively (6). In Gram-positive species, Small Alarmone Synthase (SAS) proteins, which have homology to the synthase domains of RSH proteins (4), also produce (p)ppGpp (7) in a non-ribosome dependent fashion.

Extensive amino acid contacts with the ribosome mediate the regulation of RSH-dependent (p)ppGpp synthesis in response to amino acid availability (8–10). Additionally, the RSH proteins *E. coli* RelA (11) and *B. subtilis* Rel (10) are allosterically activated by pppGpp binding. Allostery is also observed with the *B. subtilis* SAS SasB, where mutation of a single residue prevents allosteric stimulation *in vitro* (12, 13). Introduction of this mutation into the chromosomal copy of *sasB* mimics a Δ*sasB* mutation in the context of the regulation of protein synthesis (13). However, the overall role of allostery in the regulation of (p)ppGpp metabolism *in vivo* is not well characterized.

Many RSH enzymes also exhibit hydrolase activity that converts pppGpp and ppGpp into GTP and GDP, respectively (6). By comparison with (p)ppGpp synthesis, regulation of RSH-dependent (p)ppGpp hydrolysis is less well understood. In *E. coli* the RSH protein SpoT serves as the primary hydrolase and strains expressing a hydrolysis-deficient *spoT* mutant display increased (p)ppGpp abundance during a diauxic shift (14). SpoT hydrolysis activity is regulated by protein-protein interactions with various proteins (15, 16) PMID: 37926282. Most RSH proteins contain a small molecule binding domain (ACT) and the binding of branched chain amino acids to ACT can stimulate (p)ppGpp hydrolysis in *R. capsulatus*, but the generality of this mechanism is not known (17). In addition, some bacteria contain so-called Small Alarmone Hydrolase (SAH) enzymes capable of hydrolyzing (p)ppGpp (4) but their regulation is not well understood.

RSH enzymes, particularly those of the RelA/SpotT/Rel class, exhibit a functional coupling of synthase and hydrolytic activities, mediated, at least in part, by pppGpp-dependent allosteric regulation and also fundamentally by the aminoacylation state of the tRNA in the ribosome A site. (18–20). Thus, under conditions where the population of tRNAs is partially aminoacylated (21), RSH enzymes would likely oscillate between synthase and hydrolase activities resulting in changes in (p)ppGpp abundance during growth. Such a mechanism could underlie the proposed ability of (p)ppGpp to act as a reversible brake on growth (22). In fact, during strong heat shock, the abundance of (p)ppGpp increases quickly (within 2 min) but is substantially dissipated by 15 min (3). Existing methods of measuring (p)ppGpp abundance including radiolabeling of cells with ^32^P-ATP under low phosphate conditions followed by analysis of cell extracts by thin layer chromatography (TLC) or liquid chromatography/mass spectrometry (LC/MS) analysis of lysates (e.g., (23–26)) are relatively cumbersome and are not optimal techniques for conditions such as growth that may require both temporal sensitivity and physiological robustness.

Here, we use a (p)ppGpp-sensitive riboswitch (27) in conjunction with firefly luciferase (luc), an unstable reporter with a half-life in *B. subtilis* of ∼5 min (28), to serve as a dynamic sensor of (p)ppGpp abundance. We use this sensor to directly demonstrate the roles of the hydrolytic activity of the RSH protein Rel and allosteric regulatory sites in both Rel and a SAS protein in governing changes in (p)ppGpp abundance during physiological contexts including nutrient exhaustion and amino acid downshifts. In addition, we demonstrate that (p)ppGpp synthesis closely coincides with the expression of genes under (p)ppGpp control, indicating that this regulation is rapid and likely direct.

## Results

We placed a sequence corresponding to the (p)ppGpp-sensitive riboswitch from the promoter of *Desulfitobacterium hafniense ilvE* (27) between an inducible promoter (P*_hyperspank_*) and the *luc* gene encoding firefly luciferase (luciferin 4-monooxygenase) (RsFluc; Fig. 1A). This construct was integrated at the *sacA* locus and luminescence measurements were performed during growth in S7/glucose defined medium in a microplate reader. Raw luminescence readings were normalized to OD_600_ (RLU/OD) (Fig. 1B, blue). We observed two waves of luminescence, one just after initiation of growth and then a second larger one during early transition phase, near the time of departure from the most rapid period of growth. While the time and magnitude of the second wave (∼220 min) was robust across numerous biological replicates, the first wave was much more variable, possibly as a result of the proximity to the culture dilution performed at the initiation of the experiment. Both waves were dependent on the presence of the riboswitch in the reporter construct as a reporter (RsFluc^0^) lacking the intervening riboswitch sequence exhibited greatly reduced luminescence (Fig. S1A). The residual response suggests that (p)ppGpp may be also stimulating transcription initiation from P*_hyperspank_*.

**Fig. 1.**
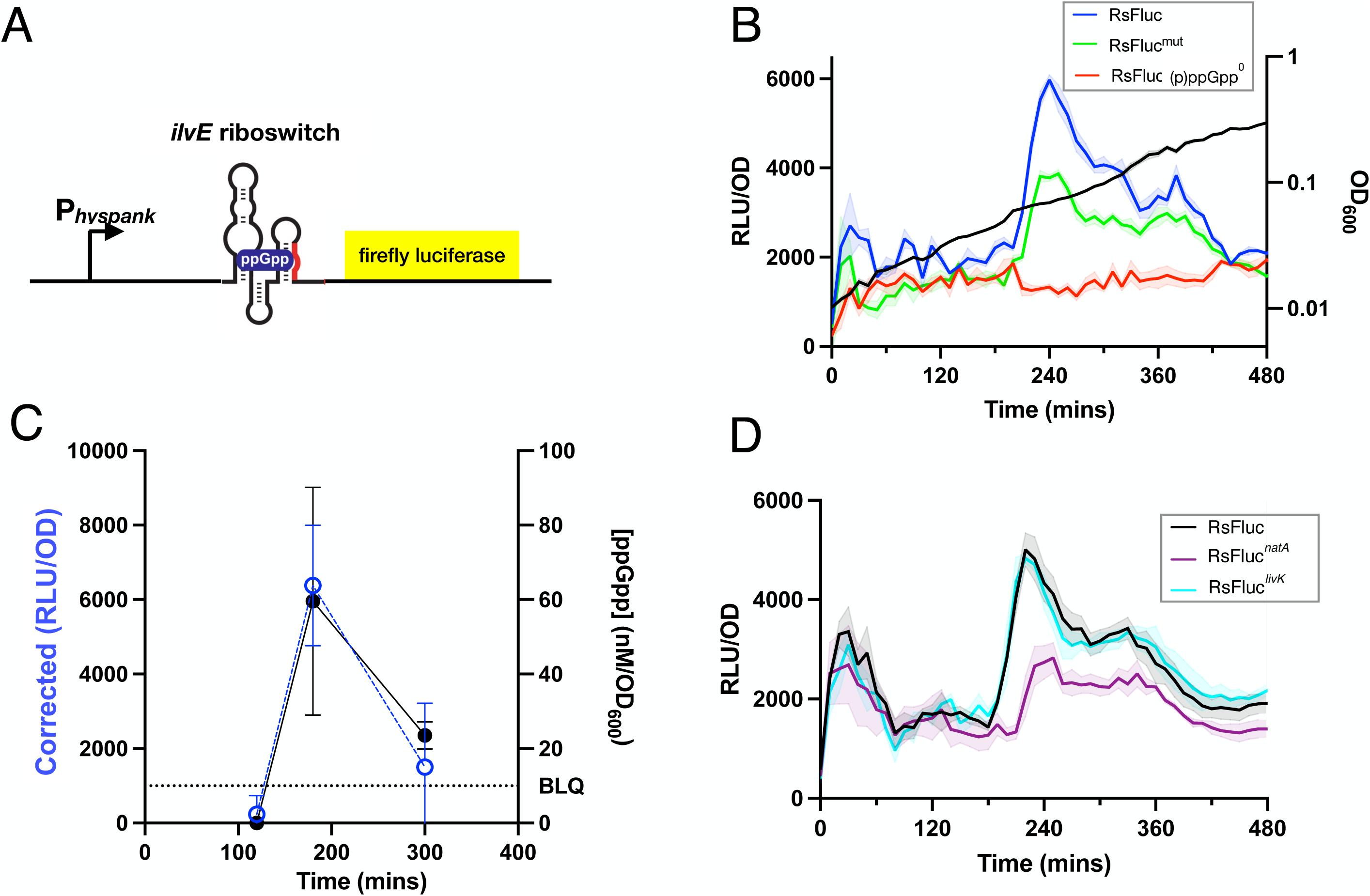
RsFluc (p)ppGpp-sensitive firefly luciferase reporter. **A**, schematic of the (p)ppGpp reporter RsFluc depicting the inducible promoter P_hyperspank_ fused to the (p)ppGpp-senstive riboswitch of *D. hafniense ilvE* followed by the gene encoding firefly luciferase. **B,** luminescence (RLU/OD_600_) of RsFluc (blue; JDB4496), mutant RsFluc^mut^ (green; JDB4631) in wildtype backgrounds and RsFluc in a Δ*relA*Δ*sasA*Δ*sasB* background (red; JDB4512), **C)** ppGpp concentration determined by HPLC-MS (black closed circles) and corresponding reporter luminescence corrected for (p)ppGpp^0^ luminescence by subtraction (blue open circles). **D,** luminescence per OD_600_ of RsFluc (black), RsFluc*^natA^* (purple, JDB4730) and RsFluc*^livK^*(cyan, JDB4731). Shown are representative examples of at least three biological replicates, each comprising three technical replicates.

An RsFluc-expressing strain carrying deletions in the gene encoding all three *B. subtilis* (p)ppGpp synthases (*relA*, *sasA*, *sasB*; (p)ppGpp^0^) exhibited a significantly attenuated luminescence (Fig. 1B, red), consistent with the expectation that the signal reflects (p)ppGpp abundance. In addition, we constructed mutant RsFluc reporters containing two previously identified riboswitch mutations, one that decreases sensitivity to (p)ppGpp as the binding pocket has been disrupted (M9) and a second that strengthens the terminator (M11), thereby increasing the specificity of the response (27). Reporters containing either mutation exhibited reduced luminescence (Fig. S1B, C) and a reporter containing both (RsFluc^mut^) was substantially attenuated (Fig. 1B, green). As these mutations are not known to completely prevent (p)ppGpp binding, consistently, this effect is less strong than in the (p)ppGpp^0^ background.

As a further confirmation of the reporter, we measured (p)ppGpp by LC/MS at points in the growth curve with different RsFluc activities. Values were normalized with respect to OD_600_ and to a parallel measurement in a (p)ppGpp^0^ strain, resulting in lower (p)ppGpp before and after the time of the luminescence peak (Fig. 1C). Calculation of the absolute concentration of (p)ppGpp yielded an estimate of ∼60 nM/OD_600_ (Fig. S2A). Of note, we observed only modest declines in GTP (Fig. S2B). Taken together, these results and those described above demonstrate that RsFluc activity reflects changes in (p)ppGpp abundance.

As noted above, (p)ppGpp refers collectively to ppGpp and pppGpp. The *D. hafniense ilvE* riboswitch has no *in vitro* preference for binding pppGpp or ppGpp (29), suggesting that RsFluc is equally sensitive to both molecules *in vivo*. Aptamers with preference for pppGpp or ppGpp have been identified (29). For example, the *Clostridiales bacterium natA* aptamer exhibits a ∼10-fold preference for pppGpp as compared to ppGpp whereas the *Oxobacter pfennigii livK* aptamer exhibits a ∼6-fold preference for ppGpp (29). We constructed versions of the RsFluc reporter containing these aptamers in place of *D. hafniense ilvE*, RsFluc^natA^ and RsFluc^livK^, respectively. The similarity between RsFluc and the ppGpp sensitive reporter RsFluc^livK^ (Fig. 1D) is consistent with previous observations that the abundance of ppGpp > pppGpp (30). In addition, the pppGpp sensitive reporter RsFluc^natA^ exhibited a delayed expression in comparison to the other reporters (Fig. 1D).

We then characterized the effect on RsFluc activity of perturbations that stimulate *in vivo* (p)ppGpp synthesis. As uncharged tRNAs activate RelA-dependent (p)ppGpp synthesis (5), many such perturbations affect aminoacyl-tRNA charging. One example is a culture grown in amino acid-replete medium shifted to a medium lacking amino acids, which is a transition that reduces aminoacylation (31) and stimulates (p)ppGpp synthesis (32). Consistently, a spike in RsFluc activity occurred shortly (∼30 min) after the down-shift (Fig. 2A, black) that is attenuated in strain expressing the RsFluc^mut^ reporter. The timing of this response illuminates the temporal relationship between our reporter and (p)ppGpp abundance. Amino acid starvation results in rapid diminution of tRNA charging (∼5 min, (33)) and increased ppGpp synthesis (∼10-15 min (34)). The difference between these values and that which we observe with RsFluc suggests that activation of the riboswitch and subsequent expression of the firefly luciferase together take <20 min.

**Fig. 2.**
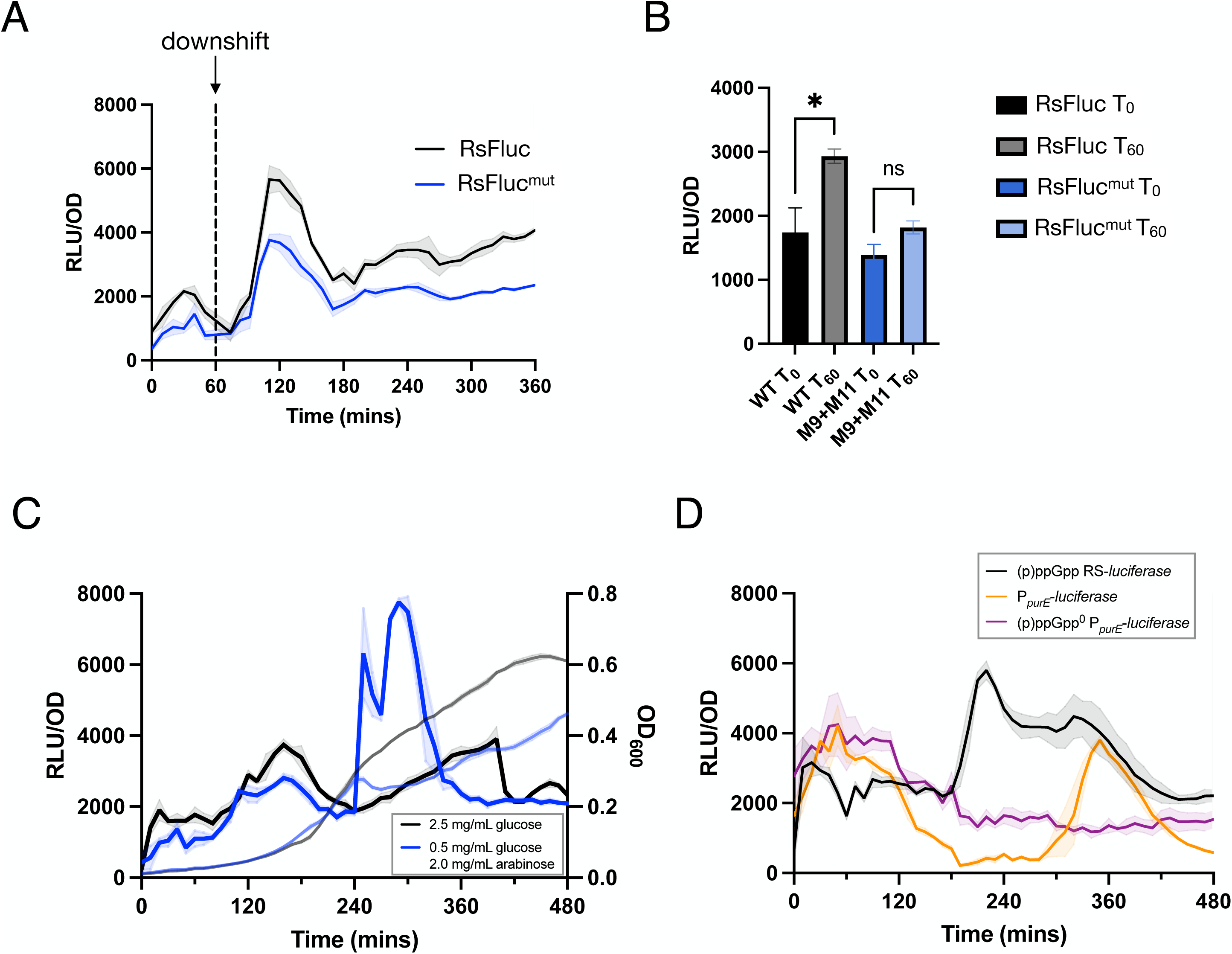
RsFluc reporter response to stimulation of (p)ppGpp synthesis. Luminescence (RLU/OD_600_) of: **A,** RsFluc (black; JDB4496) and RsFluc^mut^ (blue; JDB4631) after amino acid downshift at T_60_ (dashed line); **B,** RsFluc (black) and RsFluc^mut^ (blue) after 0 and 60 mins of 50 ng/mL mupirocin treatment. Significance was determined by two-tailed t test: p-value <.05 for RsFluc, no significant difference for RsFluc^mut^; **C,** RsFluc grown in defined minimal media containing either 2.5 mg/mL of glucose (black) or 0.5 mg/mL glucose and 2.0 mg/mL arabinose (blue). Respective OD_600_ measurements are shown in grey and light blue, respectively, and **D)** RsFluc (black) and a P*_purE_* luciferase promoter fusion in wildtype (orange; JDB4803) and (p)ppGpp^0^ (purple; JDB4804) backgrounds. Shown are representative examples of at least three biological replicates, each comprising three technical replicates.

Inhibitors of tRNA synthetases serve as an alternative strategy to modulate tRNA charging. For example, mupirocin, an inhibitor of isoleucyl-tRNA synthetase (35) stimulates (p)ppGpp synthesis (36). Consistently, RsFluc-expressing cells treated with mupirocin exhibited a significant increase in luciferase production not observed in cells expressing the (p)ppGpp-insensitive RsFluc^mut^ reporter, indicating that the effect was specific to changes in (p)ppGpp abundance (Fig. 2B). We investigated the mupirocin concentration causing the maximal stimulation of RsFluc activity, observing, surprisingly, that 100 ng/ml was less effective than 50 ng/ml (Fig. S3A). The severe diminution of growth in the presence of 100 ng/ml mupirocin (Fig. S3B) suggests at this concentration, the mupirocin is having physiological effects beyond (p)ppGpp synthesis such as direct inhibition of protein synthesis.

Both direct (^32^P-radiolabeling) (14) and indirect (transcriptional profiling) (37) assays observe (p)ppGpp synthesis during a diauxic shift when bacteria adapt to grow on a secondary carbon source following glucose exhaustion. We measured RsFluc activity during a diauxic shift from glucose to arabinose as the only carbon sources (e.g., without amino acid supplementation) following a protocol (28) based on the original Monod *B. subtilis* observations (38). The transient flattening of the growth curve in the culture grown in 0.5 mg/ml glucose & 2.0 mg/ml arabinose is characteristic of a diauxic shift and is accompanied by prominent spike in RsFluc activity at ∼240 min indicative of a substantial increase in (p)ppGpp abundance (Fig. 2C, blue). In contrast, growth in 2.5 mg/ml glucose (black) did not result in substantial changes in either growth or luciferase.

As an additional confirmation that RsFluc activity reflects (p)ppGpp abundance, we monitored expression of a firefly luciferase fusion to the *purE* promoter that is sensitive to (p)ppGpp levels (39). Specifically, (p)ppGpp enhances PurR DNA binding, thereby increasing repression of *purE* transcription. Consistent with previous observations (39), P*_purE_*-luciferase activity was affected by the absence of (p)ppGpp (Fig. 2D, compare orange and purple). We observed lowest P*_purE_*-luciferase (orange) activity at approximately the same time as RsFluc (black) activity increased (∼180 min) and a later rise in P*_purE_* activity when RsFluc was diminishing. Taken together, these data support the interpretation that RsFluc activity reflects (p)ppGpp abundance.

We then investigated the source of the (p)ppGpp responsible for the RsFluc signal. *B. subtilis* RelA, SasA (RelP), and SasB (RelQ) are the known (p)ppGpp biosynthetic enzymes (7, 40). Consistently, an RsFluc-expressing strain lacking all three proteins (Δ*relA*Δ*sasA*Δ*sasB*) exhibits substantially reduced luminescence compared to the wild type parent (Fig. 1B). We investigated the contributions of each protein to RsFluc activity by assaying cells carrying single mutations in their respective genes under growth (Fig. 3A, see Fig. S4 for corresponding growth curve) and amino acid downshift (Fig. 3B, see Fig. S6 for corresponding growth curve). While all three mutations affected expression during growth, a strain lacking RelA synthase activity due to an inactivating point mutation (*relA-D264G)* in the synthase domain (7) exhibited the largest effect (blue, Fig. 3A), essentially indistinguishable from the triple mutant Δ*relA*Δ*sasA*Δ*sasB* strain (red). In contrast, single Δ*sasA* (green) or Δ*sasB* (orange) or the combination ΔsasAΔsasB (Fig. S5, brown) mutant strains exhibited at most only modest delays in RsFluc signal. During amino acid downshift, the *relA-D264G* strain was again severely affected, but under these conditions, the Δ*sasA*Δ*sasB* strain was similarly defective (blue, brown; Fig. 3B).

**Fig. 3.**
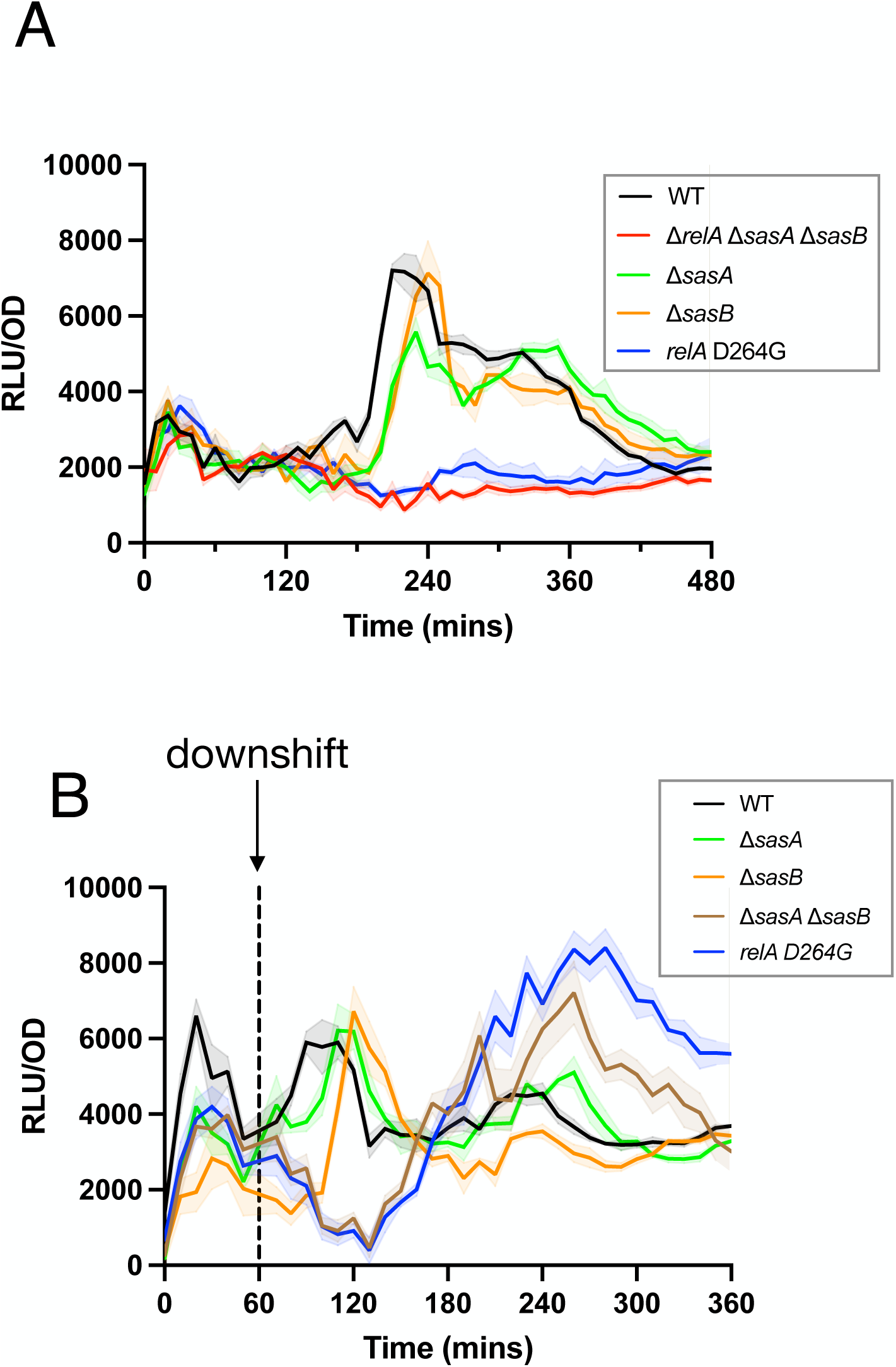
Relative contributions of (p)ppGpp synthases to dynamics of RsFluc activity. Luminescence (RLU/OD_600_) of: **A,** RsFluc in wildtype (black, JDB4496), Δ*relA*Δ*sasA*Δ*sasB* (red, JDB4512), Δ*sasA* (green, JDB4515), Δ*sasB* (orange, JDB4516), and *relA-D264G* (blue, JDB4741) backgrounds and **B),** wildtype (black), Δ*sasA* (green), Δ*sasB* (orange), Δ*sasA*Δ*sasB* (brown, JDB4511), and *relA-D264G* (blue) strains after amino acid downshift at T_60_ (dashed line). Shown are representative examples of at least three biological replicates, each comprising three technical replicates.

RsFluc activity in the Δ*sasB* strain was strikingly different as compared to the wildtype parent in the context of nutrient downshift (Fig. 3B). What underlies this effect? *In vitro*, SasB is activated allosterically by pppGpp produced by Rel (12). SasB Phe-42 is a key residue in this regulation and a SasB mutant protein carrying an F42A substitution is no longer allosterically activated *in vitro* by pppGpp (12). RsFluc activity of a strain carrying a *sasB-F42A* allele in the chromosome (13) was delayed during growth (Fig. 4A, blue) and nutrient downshift (Fig 4B, blue). Thus, this delay is the result of the absence of stimulation of SasB. Rel itself is subject to allosteric regulation *in vitro* by pppGpp (10), and Tyr-200 is important for this effect (18). Introduction of a Rel Y200A mutation into the chromosomal copy of *rel* delayed the increase in RsFluc activity during growth as compared to the wildtype parent strain (Fig. 4A, purple). During a nutrient downshift (as in Fig. 3B), this mutation also delayed increased RsFluc activity (Fig. 4B, purple). Importantly, the magnitude of the increase was not substantially affected, indicating that Rel Y200A likely does not impair synthase activity. Thus, taken together, these observations provide the first evidence of the *in vivo* function of allosteric regulation in (p)ppGpp metabolism.

**Fig. 4.**
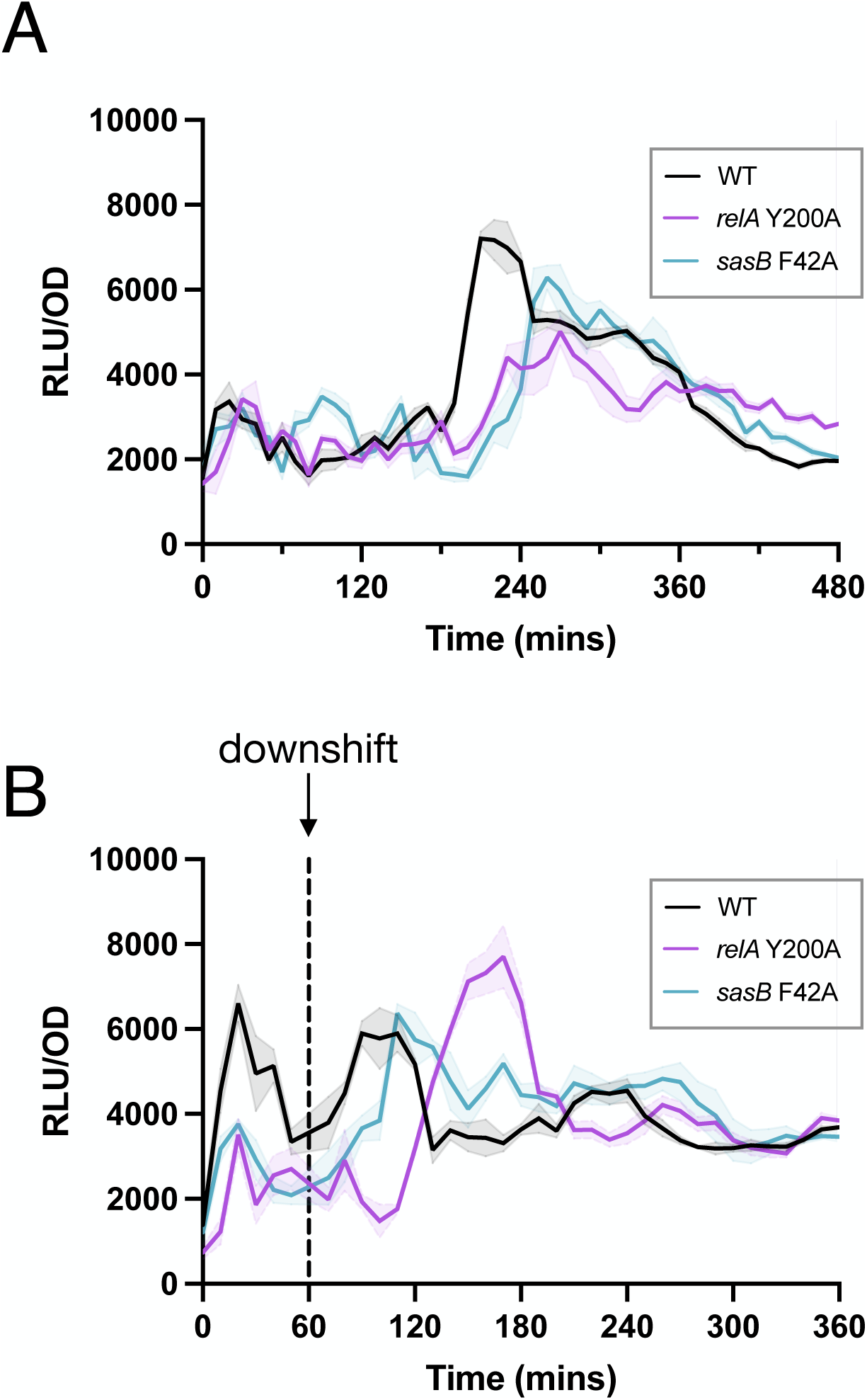
RsFluc activity coordinated by allosteric network. Luminescence (RLU/OD_600_) of: **A,** RsFluc in wildtype (black, JDB4496), *relA-Y200A* (fuschia, JDB4528), and *sasB-F42A* (blue, JDB4711) strains and **B,** RsFluc in wildtype (black), *relA-Y200A* (fuschia), and *sasB-F42A* (blue) strains after amino acid downshift (dashed line). Shown are representative examples of at least three biological replicates, each comprising three technical replicates

Regulation of RSH enzymes is not limited to direct effects on their biosynthetic activity as many RSH enzymes including Rel also hydrolyze (p)ppGpp, thereby indirectly antagonizing the synthetic activity. Investigating the contribution of Rel-dependent (p)ppGpp hydrolysis to RsFluc expression is complicated by its essentiality in the presence of any (p)ppGpp synthases, including Rel itself (40). Inspired by a previous study (3), we developed a system allowing transient expression of a RelA mutant lacking hydrolytic activity (D78A; (20)) under control of the P*_liaI_* bacitracin-inducible promoter (41) (Fig. 5A). We first confirmed that expression of an inducible wildtype *rel* allele complemented RsFluc activity in a ppGpp^0^ background (compare blue and gray; Fig. 5B). Expression of a Rel-D78A mutant protein slowed growth (Fig. S9), consistent with increased (p)ppGpp abundance, and produced a broadening of the increased RsFluc activity before and after the peaks seen with expression of wildtype RelA (Fig. 5B, red). Another reported (p)ppGpp hydrolase is NahA (42), but a Δ*nahA* mutation does not affect RsFluc (Fig. S10). Thus, Rel is the primary hydrolase activity affecting the dynamics of (p)ppGpp abundance under our experimental conditions.

**Fig. 5.**
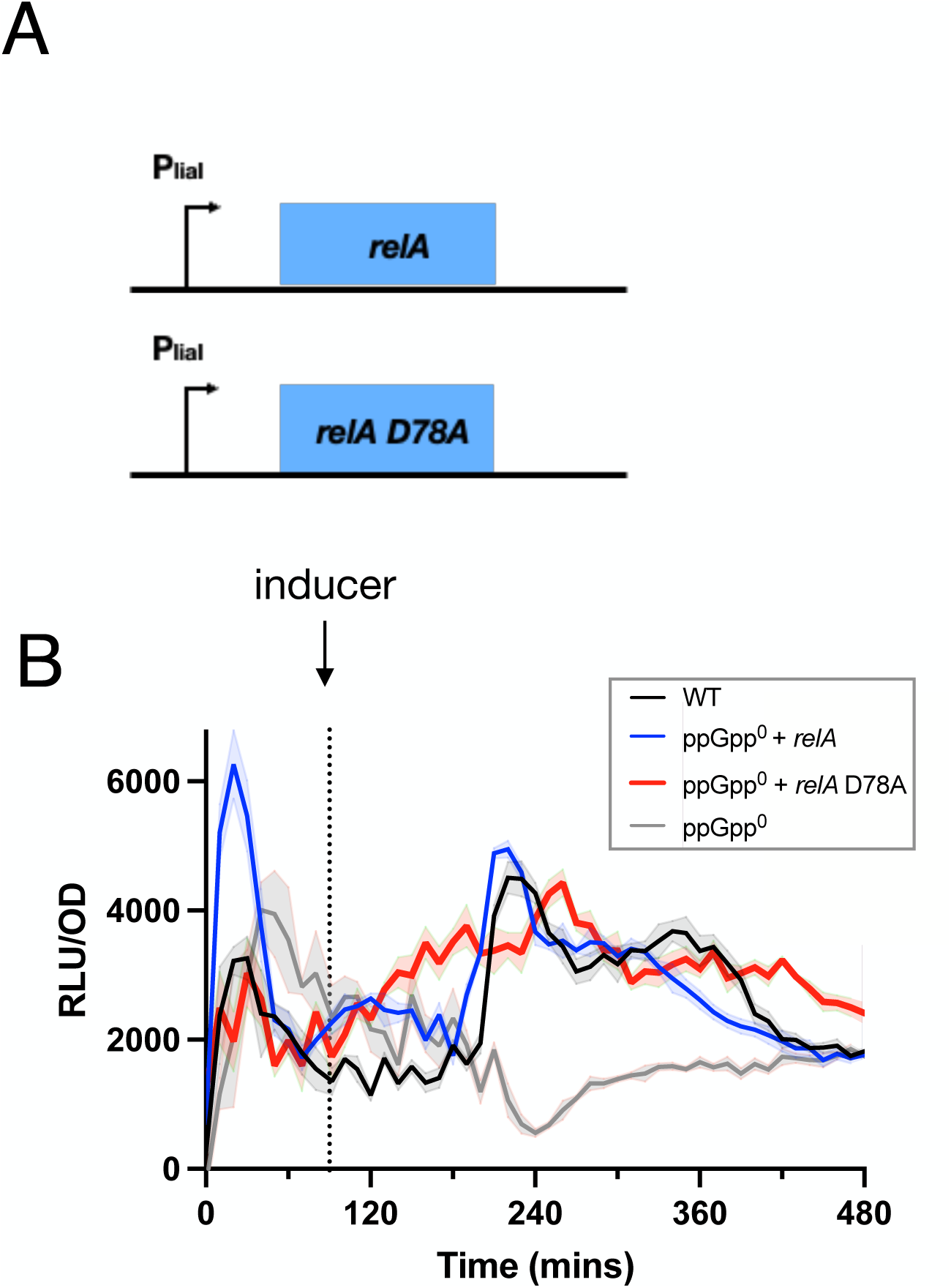
RelA hydrolase activity contributes to dynamics of RsFluc activity. **A**, schematic of bacitracin-inducible *relA* constructs. **B)** luminescence (RLU/OD_600_) of RsFluc in wildtype (black, JDB4496), Δ*relA*Δ*sasA*Δ*sasB* with P*_liaI_*-*relA* (blue, JDB4675), Δ*relA*Δ*sasA*Δ*sasB* with P*_liaI_*-*relA-D78A* (red, JDB4676) and Δ*relA*Δ*sasA*Δ*sasB* (gray, JDB4512) strains. RelA expression induced by 5 µg/mL bacitracin at T_90_ (dashed line). Shown is a representative example of at least three biological replicates, each comprising three technical replicates.

The importance of Rel (p)ppGpp synthetic activity for RsFluc expression (Fig. 4A) suggests that the primary signal responsible for stimulating (p)ppGpp synthesis is amino acid abundance. To test this hypothesis, we examined RsFluc activity as a function of amino acid concentration in the growth medium. Consistently, increasing concentrations of Casamino acids (CAA) delay maximal RsFluc activity (Fig. 6A) with minimal effects on growth. In the absence of added amino acids, RsFluc activity was substantially attenuated, especially following departure from exponential growth (Fig. 6B). This activity is substantially higher than exhibited by the RsFluc^mut^ (p)ppGpp-insensitive reporter, so it likely reflects actual (p)ppGpp abundance.

**Fig. 6.**
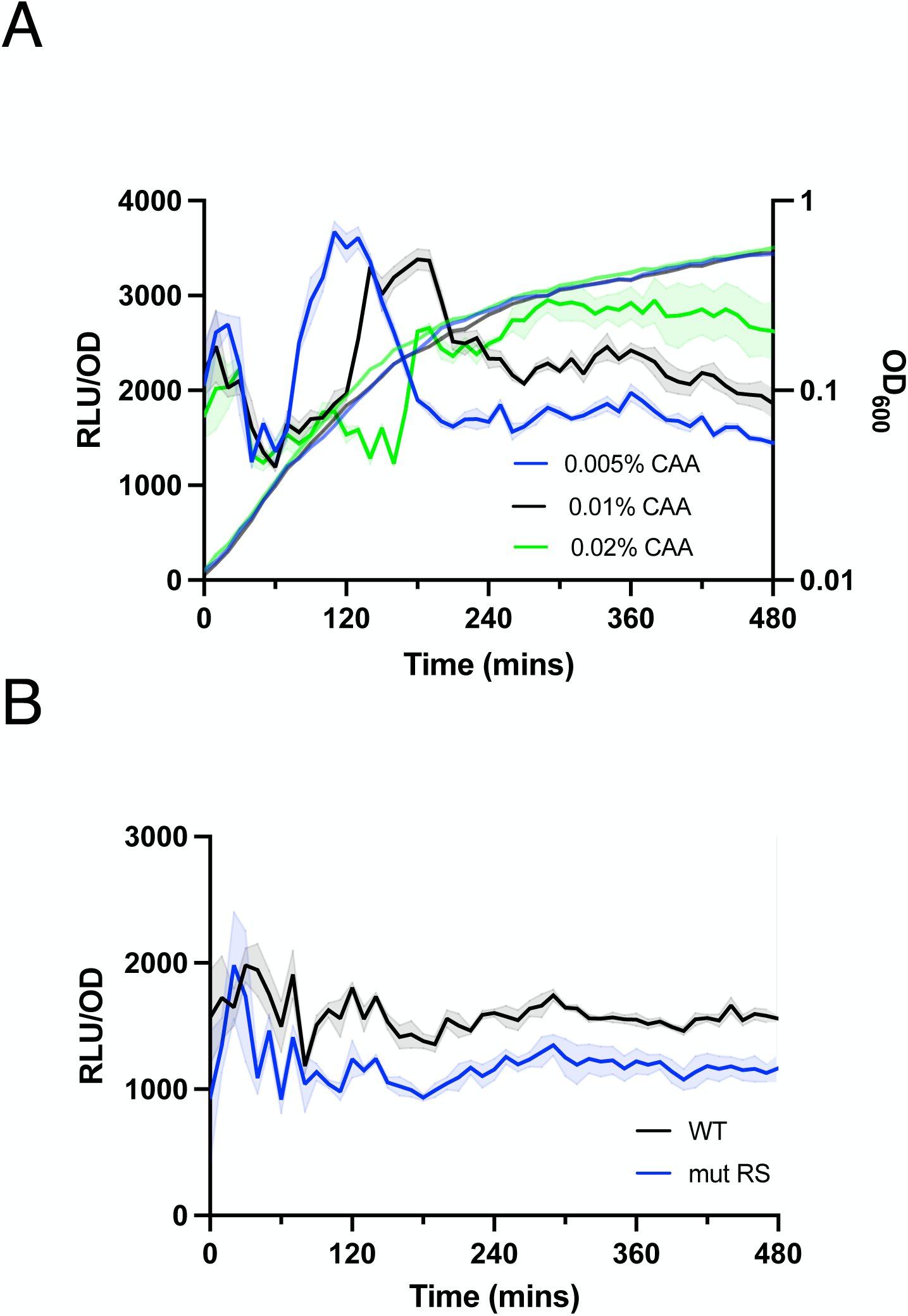
RsFluc activity and amino acid availability. Luminescence (RLU/OD_600_) of: **A,** RsFluc in prototroph (JDB4656) cultured in S7 supplemented with 0.005% (blue), 0.01% (black), and 0.02% (green) casamino acids and **B,** RsFluc (black) and RsFluc^mut^ (blue) in prototroph background without amino acid supplementation. Shown are representative examples of at least three biological replicates, each comprising three technical replicates

We finally investigated how changes in RsFluc activity correlate with the appearance of known (p)ppGpp-dependent physiological phenomena in *B. subtilis* including gene activation (43) and protein synthesis attenuation (44). Specifically, induction of the stringent response by arginine hydroxamate treatment leads to (p)ppGpp-dependent expression of a number of amino acid biosynthetic genes (43). We constructed firefly luciferase reporters of the promoters of three of these genes (*P_serA_-luc, P_ilvB_-luc, P_metE_-luc*) and compared their temporal dynamics with RsFluc. All reporters exhibited luciferase activity with very similar temporal dynamics to that of RsFluc (Fig. 7A). Consistent with the previous study, presence of a *relA-D264G* mutation substantially attenuated all reporters (Fig. 7B) (43).

**Fig. 7.**
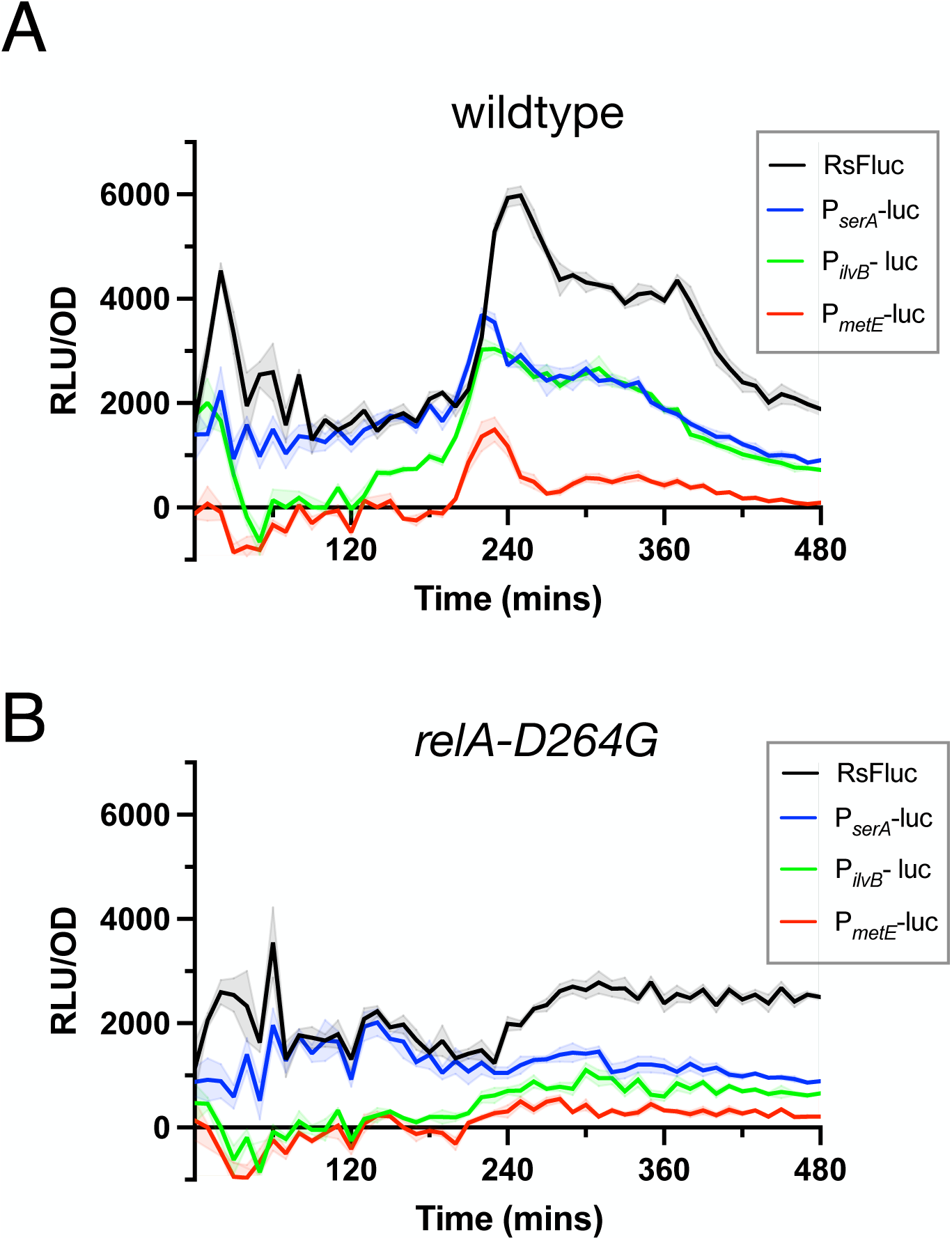
Correlation of amino acid biosynthetic gene expression and RsFluc activity. Luminescence (RLU/OD_600_) of: **A,** RsFluc (black, JDB4496), P*_serA_*-luc reporter (blue, JDB4759), P*_ilvB_*-luc reporter (green, JDB4792), and P*_metE_*-luc reporter (red, JBD4798) in the wildtype background and **B,** reporters as A in the *relA-D264G* genetic background. Shown are representative examples of at least three biological replicates, each comprising three technical replicates.

The increase in RsFluc coincides with a sharp reduction in growth rate (Fig. 8A, black). The linear correlation between protein synthesis (particularly ribosomal proteins) and growth rate (45) suggests that the increase in RsFluc is associated with reduced protein synthesis. To compare the temporal dynamics of protein synthesis and (p)ppGpp, we incorporated OPP (O-propargyl-puromycin) into cells before and during the RsFluc increase (Fig. S11, arrows “TP1”, “TP2”, respectively). Click-conjugation of OPP with a fluorophore (Fig. 8B, ‘wildtype’) and subsequent quantification revealed OPP labeling substantially declined in the interval from TP1 to TP2 (Fig. 8C). Since (p)ppGpp inhibits translation initiation (44), we asked if this change in OPP labeling was dependent on (p)ppGpp. We obtained samples of ppGpp^0^ cells at the same time points and processed them similarly (Fig. 8B, ‘ppGpp^0^’). In contrast to the wildtype cells, OPP labeling of ppGpp^0^ cells did not decline between TP1 and TP2, and in fact, it increased (Fig. 8D), consistent with the role of (p)ppGpp in attenuating protein synthesis.

**Fig. 8.**
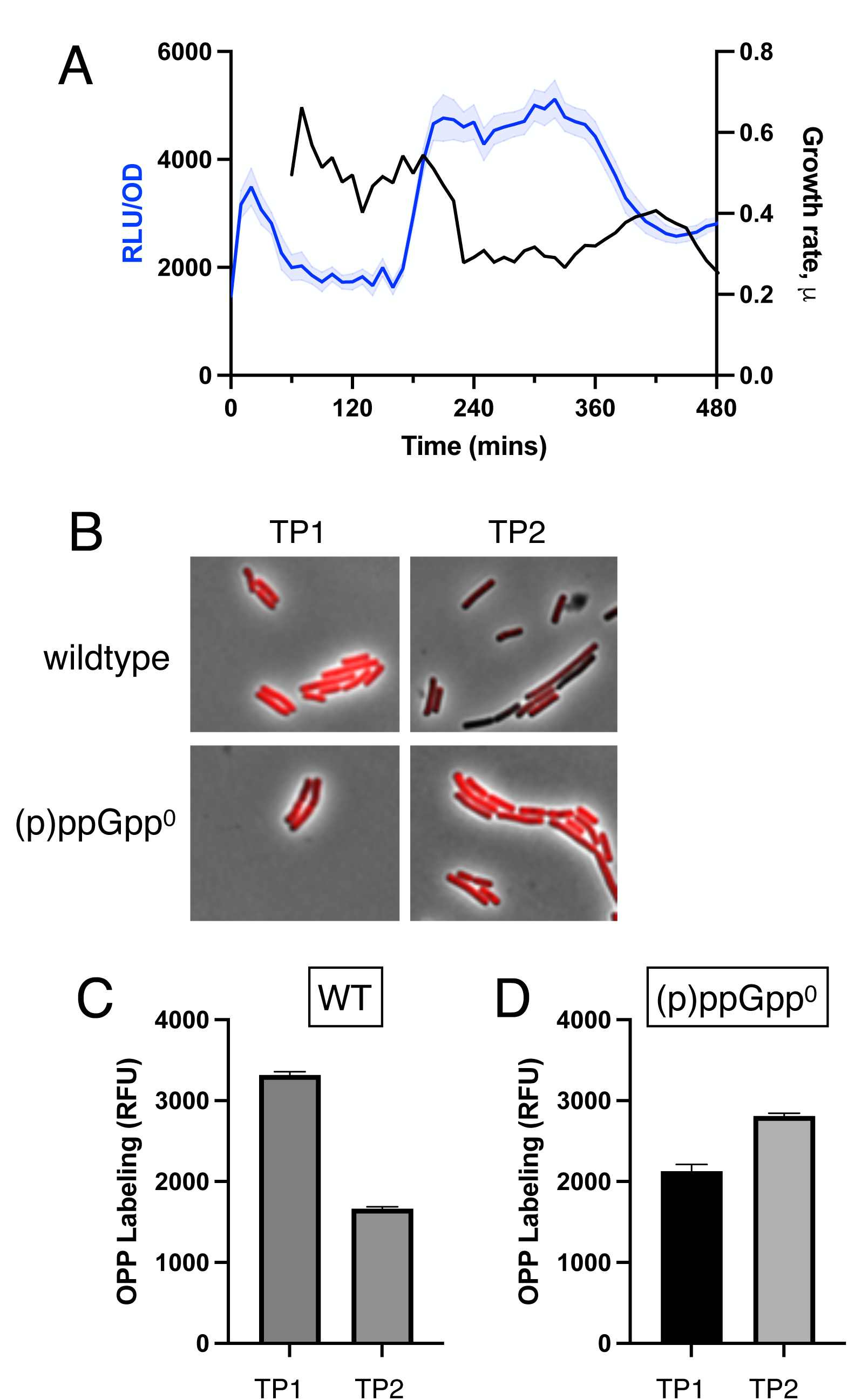
Temporal relationship between protein synthesis and RsFluc activity. **A**, luminescence (RLU/OD_600_) of RsFluc (blue, JDB4496) and growth rate (black). Growth rate/hour (μ) at a given time (t) is defined as log_2_[OD_600_(t)-OD_600_ (t_-60_)]. **B,** representative composite (fluorescent, phase) images of cells at TP1 and TP2 (see Supp Fig. 11) in either wildtype (JDB4486) or Δ*relA*Δ*sasA*Δ*sasB* (‘(p)ppGpp^0^’; JDB4512) backgrounds. **C)** and **D)** quantification of images from B.

## Discussion

Here we develop and assess RsFluc, a novel dynamic riboswitch-based reporter of (p)ppGpp metabolism in *B. subtilis*. We confirm that RsFluc accurately reflects (p)ppGpp abundance using a variety of approaches, including mutagenesis of both reporter and host. In addition, known stringent response inducers such as inhibition of tRNA aminoacylation or a nutrient downshift robustly stimulate RsFluc activity. We use RsFluc to demonstrate the *in vivo* relevance of mutations that affect allosteric activation of (p)ppGpp synthases and the involvement of Rel hydrolase activity in (p)ppGpp abundance. Together, these observations demonstrate that RsFluc will serve as an integral component of the experimental toolkit in investigating (p)ppGpp metabolism and regulation. Finally, we demonstrate the temporal dynamics of RsFluc activity correlate with the appearance of known (p)ppGpp-dependent physiological phenomena in *B. subtilis* including gene activation and protein synthesis attenuation, suggesting that these effects are likely direct.

The importance of Rel, the sensor of amino acid availability (Fig. 3A), and the effect of supplemental amino acid concentrations for RsFluc activity (Fig. 6A) are consistent with the long-held view that amino acid limitation is the key signal underlying (p)ppGpp synthesis. To further investigate this, we grew a prototrophic strain in the absence of supplemental amino acids where amino acid abundance is limited only by the availability of carbon and nitrogen. We observed no substantial fluctuations in RsFluc activity during growth (Fig. 6B) and reduced levels throughout growth. Importantly, the basal level of RsFluc activity is higher than that of RsFluc^mut^ (compare black and blue), indicating that these levels, even if low, are real. This data may be relevant to understanding the role of (p)ppGpp role in contexts sometimes referred to “basal” levels of (p)pGpp, which are defined as non-stressful growth conditions (22, 46, 47) such as motility during exponential growth (48) and the coordination of ribosome synthesis to functional tRNA abundance (21, 49).

Investigation of (p)ppGpp metabolism in both *E. coli* (50) and *B. subtilis* (51) observed a decrease in GTP abundance accompanying an increase of (p)ppGpp abundance consistent with GTP/GDP serving as the biosynthetic precursor. However, similar to *E. coli* during nutrient limitation conditions (23) or *B. subtilis* during heat shock (3), we do not observe a similar (negative) correlation (Fig. S2B). This discrepancy could be a consequence of differences in the kinetics of activation, as the mupirocin used in the earlier studies rapidly and strongly simulates (p)ppGpp abundance in <5 min. Thus, rapid synthesis of (p)ppGpp could deplete GTP/GDP pools before homeostatic mechanisms (52) are able to restore their levels. Alternatively, this discrepancy may reflect the magnitude of changes in tRNA charging: under explicit amino acid starvation, tRNA charging falls to near-zero levels on a time scale of ∼min (33), a much more substantial decrease than observed during growth phase transitions (21).

Our work confirms the previously reported (7) importance of Rel for (p)ppGpp synthesis during growth (blue; Fig. 3A). However, SasA and SasB only modestly contribute to the RsFluc dynamics during transition phase (orange, green; Fig. 3A). The lack of a substantial effect of Δ*sasB* is surprising given that it is expressed just prior to transition phase and is also a potent producer of ppGpp, at least *in vitro* or when overexpressed ectopically in *E. coli* (7). The lack of a substantial effect of Δ*sasA* is also surprising given that SasA is expressed during the transition from exponential growth (53)) similar to the peak in RsFluc activity and that SasA synthesizes ppGpp (26)). NahA may degrade a substantial fraction of the ppGpp to pGpp (42) but the absence of NahA does not have a substantial effect on RsFluc (Fig. S10). To our knowledge, previous studies have not examined the specific synthase and hydrolase dependency of (p)ppGpp accumulation during *B. subtilis* growth.

In contrast to growth, *sasA* and *sasB* have substantial effects during amino acid downshift (Fig. 3B, green, orange), with strains carrying Δ*sasA* or Δ*sasB* mutations exhibiting delayed RsFluc expression. In fact, the double Δ*sasAΔsasB* mutant strain (brown) exhibits greatly attenuated RsFluc activity, similar to the *relA-D264G* strain (blue). Both strains also exhibit severe growth defects following downshift (Fig. S6, blue, brown), suggesting that all three genes play a key role in the resumption of growth following abrupt starvation, as has been suggested for *E. coli* RelA (54). Thus, taken together, our observations provide a starting point for future investigations to understand the regulation of *B. subtilis* Rel, SasA, and SasB during regrowth.

The role of (p)ppGpp in the phenomenon of diauxie has been studied for >50 years (55). However, *E. coli* strains lacking (p)ppGpp exhibit multiple amino acid auxotrophies (56), complicating analysis in a growth medium consisting only of a single carbon source (that is, without added amino acids that can serve as a source of carbon). Similar auxotrophies also have been reported for *B. subtilis* (37, 43). The experiments presented here demonstrate that (p)ppGpp is synthesized during a diauxic shift in *B. subtilis* (Fig. 2C). Rel is the only (p)ppGpp synthase necessary for this synthesis as the RsFluc activity of a Δ*sasAΔsasB* strain is essentially the same as the wildtype parent (Fig. S5). This result demonstrates that during glucose exhaustion, the availability of amino acids necessary for charging tRNAs as sensed by Rel is the signal stimulating (p)ppGpp synthesis. Interestingly, *Lactococcus lactis* exhibits single cell heterogeneity in their ability to react to diauxie (57). Given that this phenomenon is subject to stringent response control, heterogeneity in (p)ppGpp abundance at the single cell level was proposed to underlie this phenomenon. This intriguing hypothesis awaits the development of (p)ppGpp reporters amenable to single cell analysis.

Allosteric regulation has been observed *in vitro* for *B. subtilis* (p)ppGpp synthases Rel (10) and SasB (12) (but not SasA (58)). Our analysis of strains carrying mutations in residues identified *in vitro* to be important for this regulation (RelA-Y200A (10), SasB-F42A (12) ) reveals that that allostery affects (p)ppGpp abundance *in vivo* (Fig. 4). In addition, while these strains grow equivalently to the wildtype parent (Fig. S7), both exhibit impaired recovery from nutritional downshift (Fig. S8). This is the first time that allostery has been demonstrated to mediate a specific physiological function of the (p)ppGpp metabolic network. Since strains carrying a Δ*sasB* deletion also exhibit a substantial phenotype with respect to (p)ppGpp synthesis in response to a nutritional downshift (Fig. 3B), the phenotype of a *sasB-F42A* strain with respect to recovery suggests a mechanistic explanation for this phenotype.

The synthetic activity of *B. subtilis* (p)ppGpp synthases is well understood to be regulated by interactions with the ribosome (e.g., Rel (20)) or by transcriptional regulation in the case of SasA/RelP and SasB/RelQ (53)). In contrast, the regulation of hydrolase activity, either in the context of the Rel dual synthase/hydrolase or standalone hydrolases such as NahA is less clear. A difficulty in investigating the in *vivo* function of Rel hydrolase activity results from its essentiality in the presence of any (p)ppGpp synthetic activity. Our experiments using comparison of a transiently expressed wildtype Rel with a hydrolase-dead mutant (D78A) directly demonstrate that the hydrolase activity of Rel affects the abundance of (p)ppGpp as verified by the RsFluc reporter (Fig. 5B). The relative constancy of luciferase expression in the *relA-D78A* strain as compared with the strain expressing a hydrolase-active allele, suggests that regulation of hydrolysis activity is important in generating the observed spike in ppGpp and suggests that it is not just regulation of the synthase domain which is important for determining (p)ppGpp abundance. This interpretation is consistent with a recent model of Rel function, envisioning a switch between Hydrolase-Active/Synthase-Inactive and Hydrolase-Inactive/Synthase-Active configurations (10, 20). This switch could be controlled by ribosome association, but it is not clear what would stimulate a change in this interaction under the gradual nutrient exhaustion investigated here. Alternatively, this could be a consequence of the allosteric regulation of Rel by one of its products (18), as discussed above.

Treatment with a small molecule that reduces tRNA aminoacylation, such as mupirocin, a isoleucyl-tRNA synthetase inhibitor, is a typical laboratory strategy to induce (p)ppGpp synthesis (35). We observe that mupirocin stimulates RsFluc expression (Fig. 2B) but the response is not monotonic, with 50 ng/ml mupirocin stimulating to a greater extent than 25 ng/ml but less than 100 ng/ml (Fig. S4A). Given that increased mupirocin results in decreased tRNA charging, which itself inhibits protein synthesis, 100 ng/ml mupirocin could affect protein synthesis sufficiently to interfere with expression of the luciferase reporter. The observation that 100 ng/ml mupirocin affects growth much more substantially than 50 ng/ml (Fig. S3B) is consistent with this explanation. Thus, interpretation of the effect of 100 ng/ml mupirocin is complicated since ppGpp also directly inhibits protein synthesis (59). Similar issues may be common with other molecules that stimulate (p)ppGpp synthesis by affecting tRNA charging, suggesting that careful titration may be necessary in order to properly control for indirect effects on protein synthesis (60).

The close temporal correlation between the activity of luciferase fusions to the promoters of several amino acid biosynthetic genes with RsFluc (Fig. 7A) suggests that (p)ppGpp directly activates their transcription. A fall in GTP levels as a consequence of (p)ppGpp synthesis and the subsequent activation of CodY, a known repressor of amino acid biosynthetic genes is a potential mechanism (43). However, GTP levels do not significantly fall during the time of RsFluc activity (Fig. S2B), similar to that observed when (p)ppGpp and GTP abundances were monitored during *E. coli* growth by HPLC (23). Finally, (p)ppGpp binds RNAP and in conjunction with DksA directly modulates transcription of target genes in *E. coli*, (61), but as there is no evidence for a similar interaction in *B. subtilis*, future research should investigate how such positive regulation occurs.

The use of riboswitches as a basis for physiological reporters of signaling nucleotide abundance is becoming increasingly widespread. Examples include a *B. subtilis* cyclic-di-GMP reporter (62), a *S. aureus* cyclic-di-AMP reporter (63), an *E. coli* (p)ppGpp reporter (64). These studies utilized a fluorescent reporter and observed changes in fluorescence during growth. These metabolites, especially (p)ppGpp, play important roles in the physiology of commensal and pathogenic bacteria. For example, bacteria lacking (p)ppGpp exhibit defects in virulence (e.g., *Campylobacter jejuni* (65)), *Mycobacterium tuberculosis* (66, 67)) and commensal persistence (68) and colonization (69). One issue in investigating these phenomena is that assessment of (p)ppGpp abundance in situations outside of carefully controlled physiological contexts is technically challenging. However, the ability to measure luciferase produced by bacteria during murine infection (70), suggests that it may be possible to monitor expression of the RsFluc reporter inside of a host. In addition, this reporter should facilitate the analysis of (p)ppGpp metabolism in natural isolates as well as in antibiotic-tolerant persister cells which display intriguing variability in (p)ppGpp abundance (71).

Amino acid availability plays a central role in host-microbe and microbe-microbe interactions. For example, the branched chain amino acids leucine, isoleucine, and valine are essential mammalian amino acids and as such, gut bacteria are a critical source of these molecules (72). In addition, extracellular amino acid availability/exchange plays a key role in the organization of bacterial communities (73). Intracellular amino acid availability affects extracellular amino acid availability, either as a consequence of passive and/or active transport across the membrane (74) or potentially as a result of phage lysis (75). Thus, since (p)ppGpp plays a central role in coordinating intracellular amino acid biosynthesis, (p)ppGpp metabolism likely affects these processes. The RsFluc reporter described here should greatly facilitate investigation of this regulation in physiologically complex contexts.

## Methods

### Strain construction

Strains were derived from *B. subtilis* 168 *trpC2* except as noted and are listed in Table S1. Strains were constructed by transformation using conventional methodology and where necessary, media was supplemented with either 100 µg/mL spectinomycin, 10 µg/mL kanamycin, 5 µg/mL chloramphenicol, or 1X MLS. Reporters at *sacA* were constructed using pSac-cm (76) derived plasmids (Table S2), confirmed via whole plasmid sequencing via Plasmidsaurus or Genewiz, and *sacA* integration was confirmed by assaying growth on TSS/glc and nongrowth on TSS sucrose plates. Constructs at *amyE* were constructed using pDG1730 derived plasmids, confirmed via whole plasmid sequencing, and *amyE* integration was confirmed by assaying on LB starch plates. Bacitracin-inducible constructs were designed as described previously (77), using P*_liaI_* promoter upstream of induced gene (Table S2). Marker free site-directed mutagenesis of the *B. subtilis* chromosome was done using pMINIMad2 methodology as described (78), confirming via whole plasmid sequencing and mutated locus sequence by amplification and Sanger sequencing (Genewiz).

### (p)ppGpp reporter construction

The various (p)ppGpp reporter strains were constructed by fusing their respective gBlocks (Table S3) downstream of IPTG-inducible promoter, P*_hyperspan_*_k_, and upstream of firefly luciferase gene, using a pSac-cm cloning vector via conventional restriction enzyme and Gibson assembly cloning techniques. Plasmids were sequenced verified and are listed in Table S2. The vectors were transformed into *B. subtilis*, selecting on LB Agar supplemented with 5 µg/mL chloramphenicol, and *sacA* integration was verified using TSS/glc and TSS sucrose plating.

### Luminescence measurements

Luminescence was measured in a Tecan Infinite M200 Pro instrument with continuous shaking at 37°C, taking OD_600_ and luminescence reads every 10 minutes. Cultures were grown from single colonies grown overnight on LB plates at 37°C. Single colonies were picked into 2 mL S7+CAA (1X MOPS (Teknova), 1.32 mM K_2_HPO_4_, 1% glucose, 0.1% glutamic acid, 0.01% casamino acids, 40 µg/mL L-Trp), unless otherwise stated, and grown at 37°C in a roller drum for ∼3.5 hours. OD_600_ measurements were taken to ensure colonies remained in early exponential phase (between OD_600_ 0.3-0.6). The cultures were then diluted to OD_600_=0.05 in 0.5 mL fresh S7+CAA OD_600_ 0.3-0.6). The cultures were then diluted to OD_600_=0.05 in 0.5 mL fresh S7+CAA supplemented with 4.7 mM D-luciferin (Goldbio) and 10 µM IPTG (Goldbio). Note that a spectrophotometer OD_600_ reading of 0.05 is equivalent to a plate reader reading of ∼0.01. Cultures were aliquoted in triplicate in wells amounting 150 µL each in a 96-well white-walled, flat and clear bottom plate (Greiner Bio-One). Media only cells were used for background subtraction.

### HPLC-MS nucleotide quantification

Cultures were grown from single colonies grown overnight on LB plates at 37°C. Single colonies were picked into 2 mL S7+CAA and grown at 37°C in a roller drum for ∼3.5 hours. OD_600_ measurements were taken to ensure colonies remained in early exponential phase (between OD_600_ 0.3-0.6). The cultures were back diluted to OD_600_ 0.05 in 20 mL S7+CAA supplemented with 4.7mM D-luciferin and 10µM IPTG in baffled flasks. The cultures were incubated at 37°C with shaking in a water bath. At 120, 180, and 300 min, 450 µL were sampled for measuring luminescence and OD_600_ via Tecan Infinite M200 Pro plate reader, aliquoting 150 µL in triplicate wells. Media only wells were used for background subtraction. Simultaneously, 5 mL of culture were concentrated on 0.2 µm pore (d = 0.47 mm) cellulose acetate membrane filter (Sartorius) via vacuum filtration. The filter was immediately washed with 1 mL ice cold 1M acetic acid solution containing 1 µg/mL ^15^N-ATP (Sigma) and 1 µg/mL ^15^N-GTP (Sigma) in a 50 mL conical tube, repeatedly washing the solution over the filter surface using the force of a pipette. Conical tubes were vortexed for 3-5 seconds, and the solution was transferred to 2 mL cryo-vials. Solutions were stored at −80°C. Cold acid nucleotide extractions were performed by thawing samples on ice for 60 minutes, vortexing occasionally. Samples were then re-frozen using liquid nitrogen and lyophilized for 6 hours using a VirTis Benchtop Freeze Dryer. Lyophilized samples were dissolved in 200 µl ice-cold HPLC-grade water, centrifuged at maximum speed for 30 minutes, and clear supernatant was collected for quantification via UHPLC-MS/MS.

UHPLC-MS/MS was performed using an ACQUITY Premier UPLC System coupled with a Waters XEVO TQ-S triple quadrupole mass spectrometer. UPLC was performed on a Hypercarb 2.1×50 mm porous graphitic carbon column (3 μm particle size) using a 10-90% linear gradient of solvent B (0.1% ammonium hydroxide in acetonitrile) in solvent A (0.1 ammonium acetate in water, adjusted to pH 9.5 with ammonium hydroxide) within 10 minutes and a flow rate of 0.3 ml/min. MS/MS analysis was operated in negative ionization mode and a multiple reaction monitoring (MRM) mode was adopted. MassLynx was used to quantify peak intensities.

### Amino acid downshift

After 60 min growth in the plate reader, cultures were collected in sterile microcentrifuge tubes, pelleted by centrifugation, and resuspended in equal volume of S7 lacking CAA. Cultures were returned to the plate reader, taking OD_600_ and luminescence reads every 10 minutes for remainder of assay.

### Mupirocin treatment

After 90 min growth in plate reader, mupirocin was added to the cultures at noted concentrations, and OD_600_ and luminescence measurements were resumed for remainder of assay.

### Diauxic shift

Cultures were grown in either 0.5 mg/ml glucose or a mixture of 0.5 mg/mL glucose and 2.0 mg/mL arabinose.

### OPP Labeling

Click-iT Plus OPP Protein Synthesis Assay Kit (Invitrogen) was used to label cells with OPP following manufacturer’s instructions. 1000 µL of cells at given time points were transferred to disposable glass tubes. OPP was added to a final concentration of 10 µM. OPP incorporation was performed at 37 °C on a roller drum for 20 min and all subsequent steps were done at RT. Cells were fixed by adding formaldehyde to a final concentration of 1%. Cells were fixed for 30 min, harvested by centrifugation at 15k x *g* for 3 mins, and permeabilized using 100 µL of 0.5% Triton X-100 in PBS for 15 min. Cells were labelled using 100 µL of 1X Click-iT cocktail for 20 min in the dark. Cells were harvested and washed one time using Click-iT rinse buffer and then re-suspended in 20-100 µL of PBS for imaging.

### Microscopy

Microscopy was performed on live cells immobilized on 1% agarose pads prepared with PBS. Imaging was performed using a Nikon 90i microscope with a Phase contrast objective (CFI Plan Apo Lambda DM ×100 Oil, NA 1.45), an X-Cite light source, a Hamamatsu Orca ER-AG, and the following filter cube: mCherry (ET Sputter Ex560/40 Dm585 Em630/75). Phase contrast and fluorescence images of bacterial cells immobilized on agarose pads were acquired. The image stacks were analyzed in the software Fiji with the help of the MicrobeJ plugin (79). The straighten and intensity options in the MicrobeJ plugin were used to measure the average fluorescence per pixel within each cell. Mann-Whitney statistical test was performed using MicrobeJ to ascertain the significance of the fold differences between the strains. A non-fluorescent control strain was used to subtract background and autofluorescence in each channel.

## Acknowledgements

We acknowledge the contributions of Abigail Whalen to the initial development of the riboswitch reporter and advice from other members of our laboratory. We thank Jonathan Jagodnik for helpful discussions about nucleotide-specific riboswitches and Frederico Gueiros Fihlo for comments on the manuscript. This work was supported by NIH R35GM141957 (JD) and US Army Research Office, contract W911NF2110015 (CH). MH gratefully acknowledges funding support from the Columbia University Graduate Training Program in Microbiology and Immunology (T32AI106711) and the Training in Cardiovascular Translational Research Training Grant (5T32HL120826).

---

**Fig. S1.**
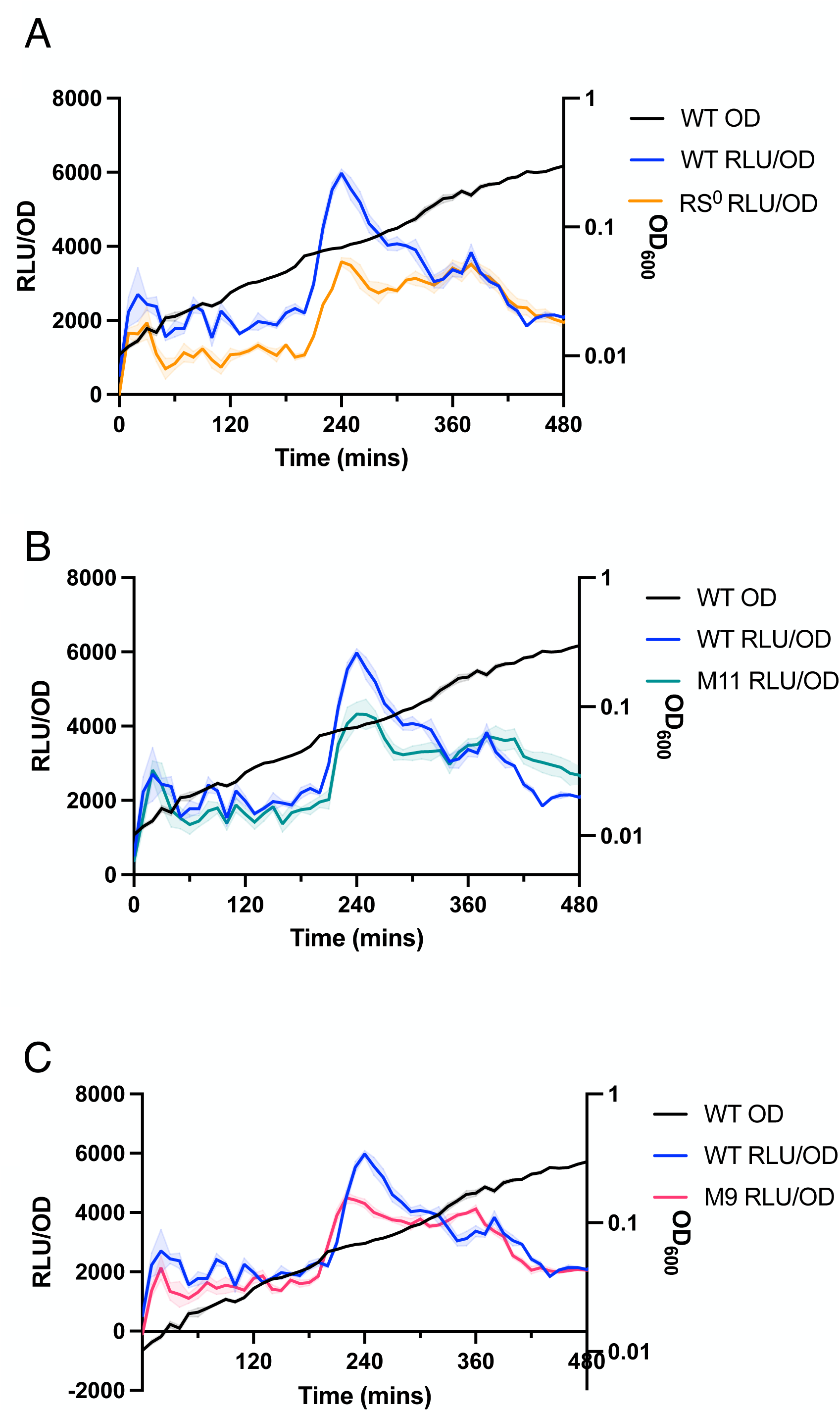
Luminescence activity of RSFluc reporter mutants. Luminescence (RLU/OD_600_) of: **A,** RsFluc (blue) and mutant RsFluc construct lacking an aptamer sequence (orange, RS^0^), **B,** RsFluc (blue) and M11 mutant RsFluc (green); and **C,** RsFluc (blue) and M9 mutant RsFluc (pink). Growth (OD_600_) (black).

**Fig. S2.**
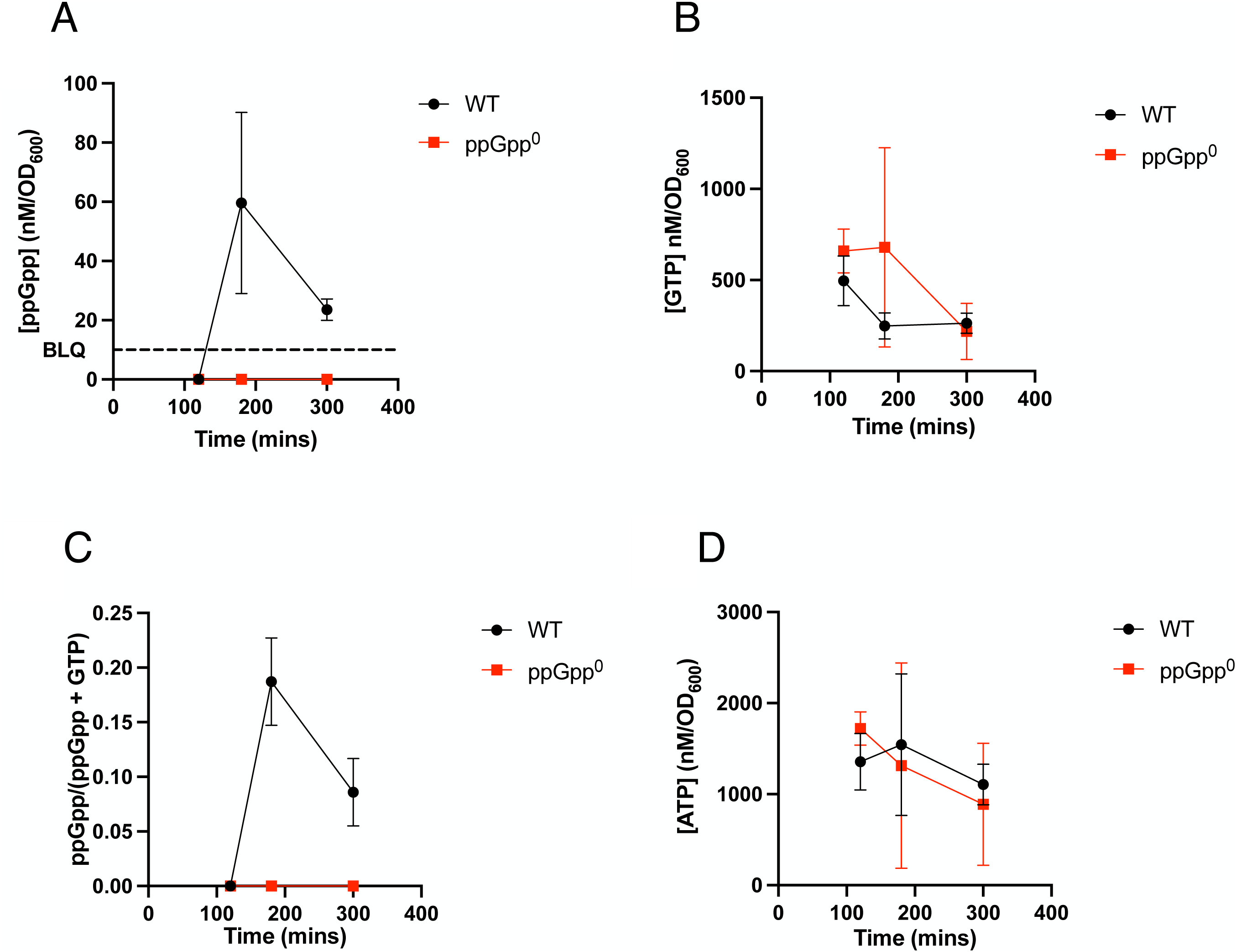
HPLC-MS quantifications of nucleotide levels sampled during growth. Nucleotide quantifications in the WT (black, JDB4496) and a (p)ppGpp^0^ background (red, JDB4512) of **A)** ppGpp, **B)** GTP, **C)** guanosine pools, and **D)** ATP, as analyzed via LC-MS collected at specified intervals.

**Fig. S3.**
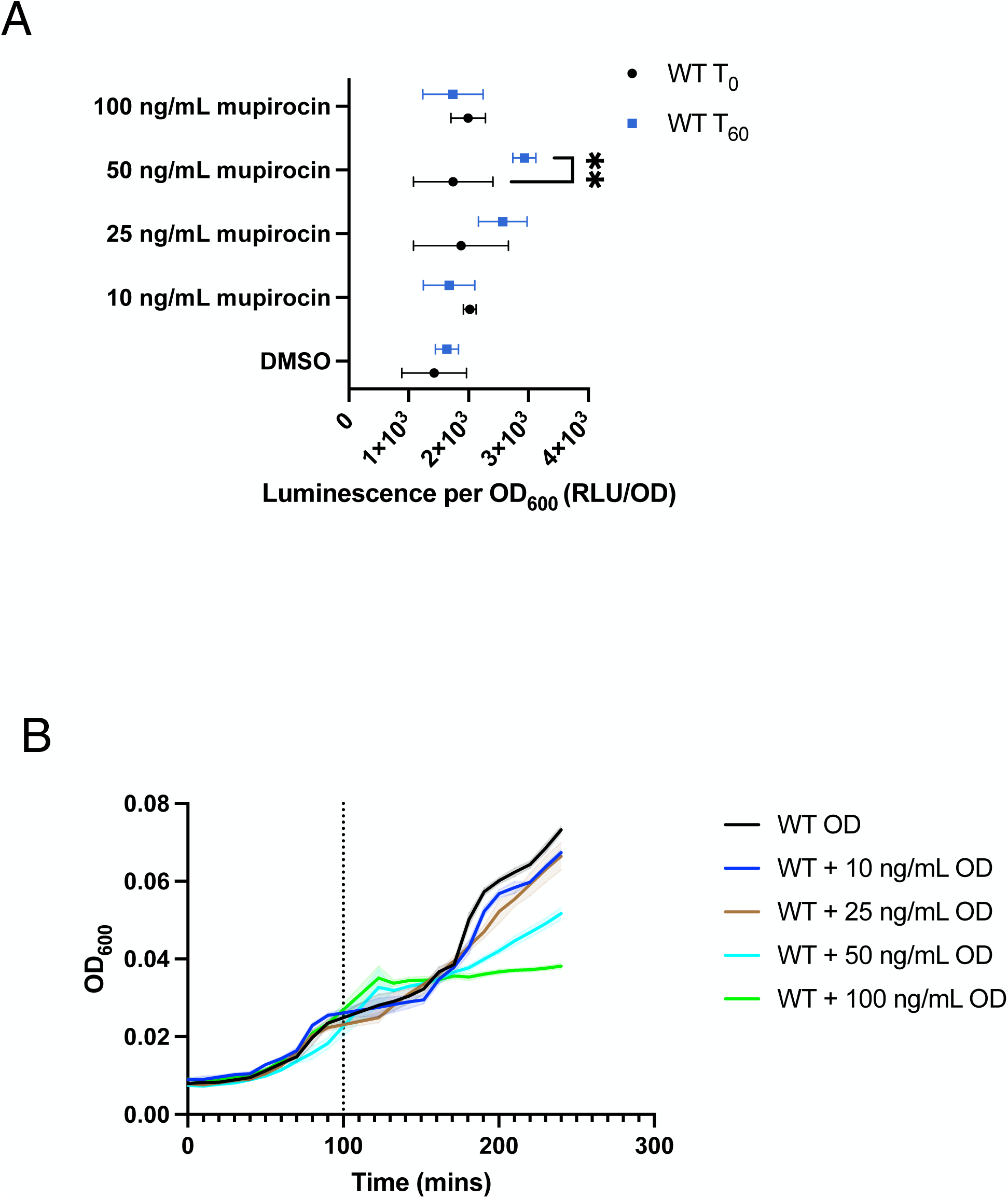
RsFluc response to mupirocin titration. **A**, luminescence (RLU/OD_600_) of RsFluc (JDB4496) at start (T_0_) of treatment (black) and after 60 mins of treatment, T_60_ (blue) with varying concentrations of mupirocin. Significance determined by two-way ANOVA with multiple comparison, comparing T_0_ and T_60_ under each treatment. 50 ng/mL mupirocin (*) had a p-value of 0.0036 whereas other treatments were not significant. **B,** growth (OD_600_) of RsFluc before and after the time of mupirocin addition (dotted line) as treated with DMSO (black), 10 µg/mL (blue), 25 µg/mL (brown), 50 µg/mL (cyan), and 100 µg/mL (green) mupirocin.

**Fig. S4.**
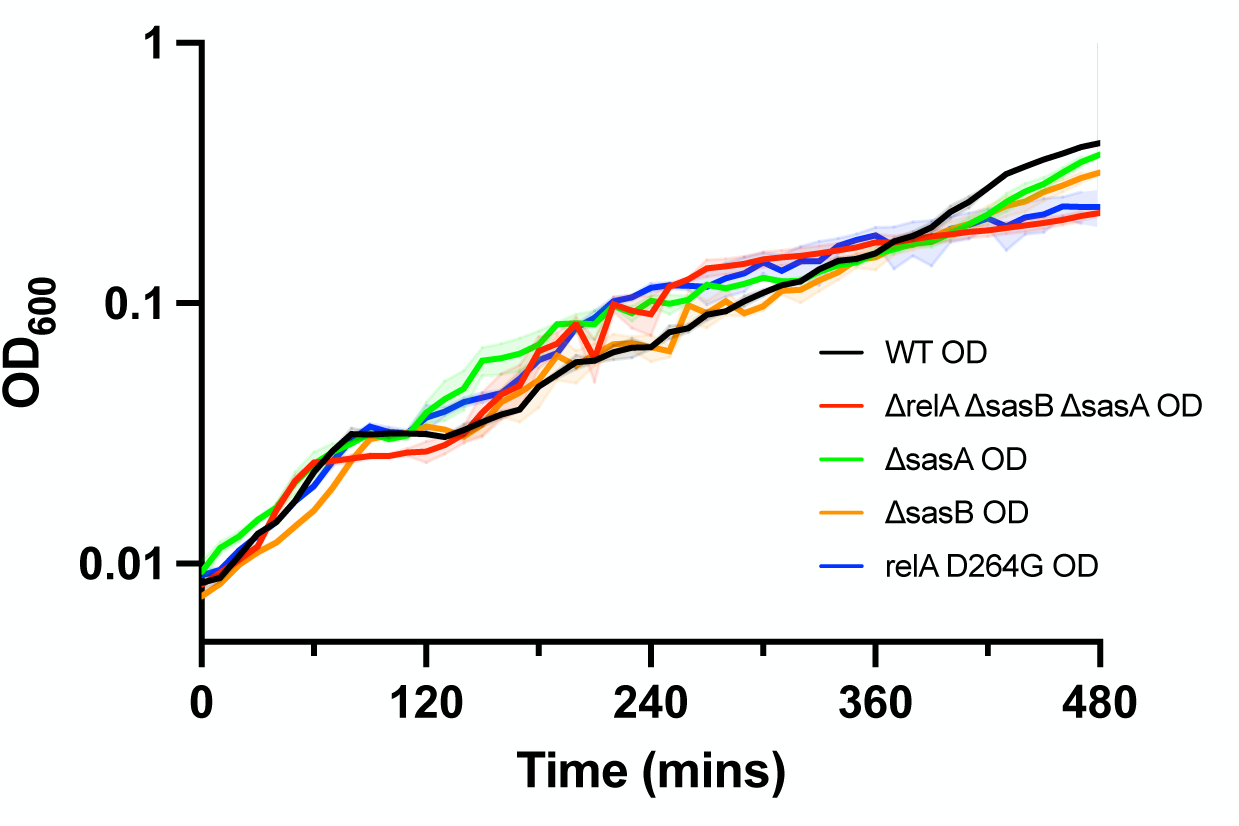
Growth curves of (p)ppGpp synthetase mutants. Growth (OD_600_) of wildtype (black, JDB4496), (p)ppGpp^0^ (red, JDB4512), Δ*sasA* (green, JDB4515), Δ*sasB* (orange, JDB4516), and *relA-D264G* (blue, JDB4741) strains.

**Fig. S5.**
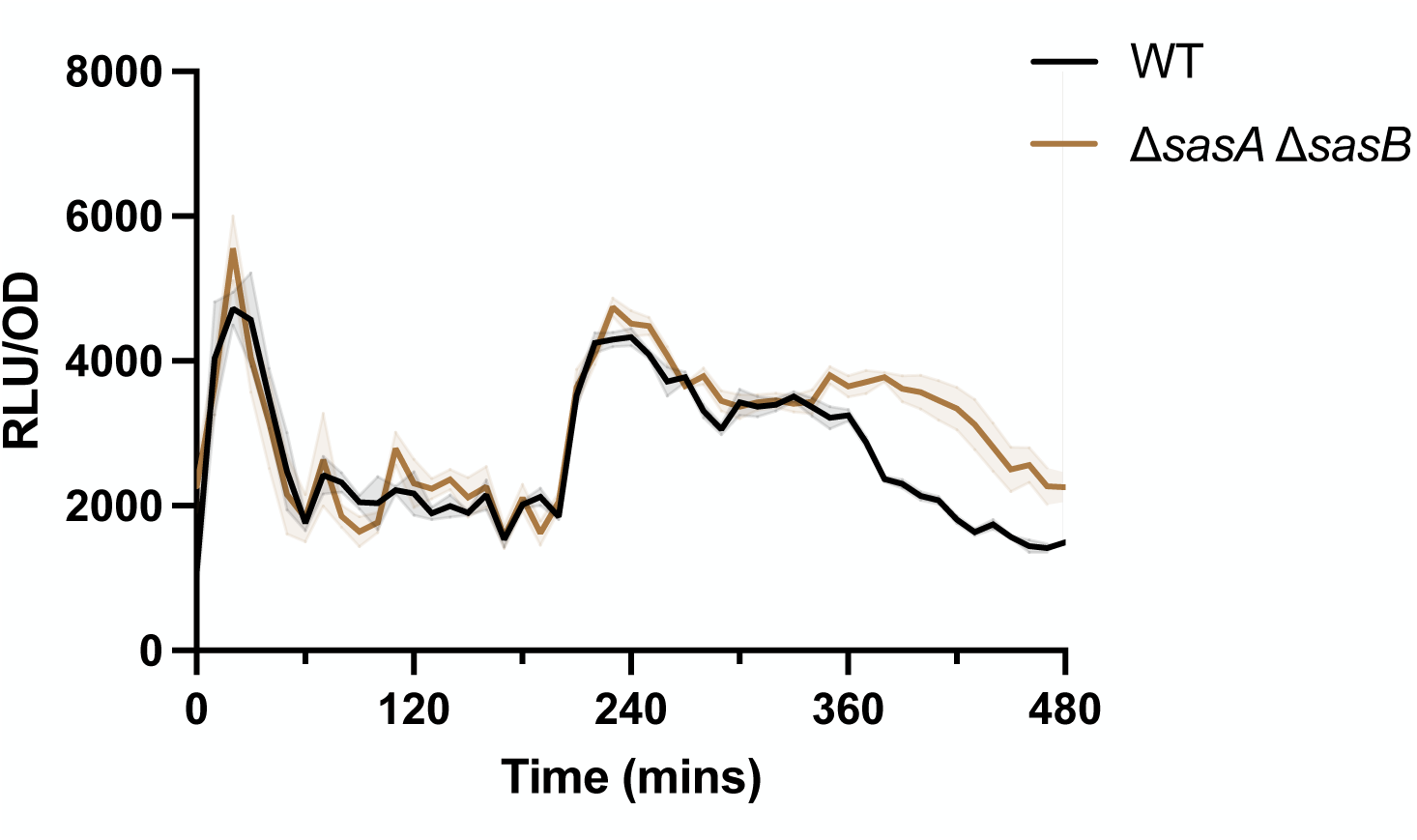
Effect of SasA/B proteins on RsFluc activity. Luminescence (RLU/OD_600_) of RsFluc in WT (black, JDB4496) and Δ*sasA*Δ*sasB* (gold, JDB4508) backgrounds.

**Fig. S6.**
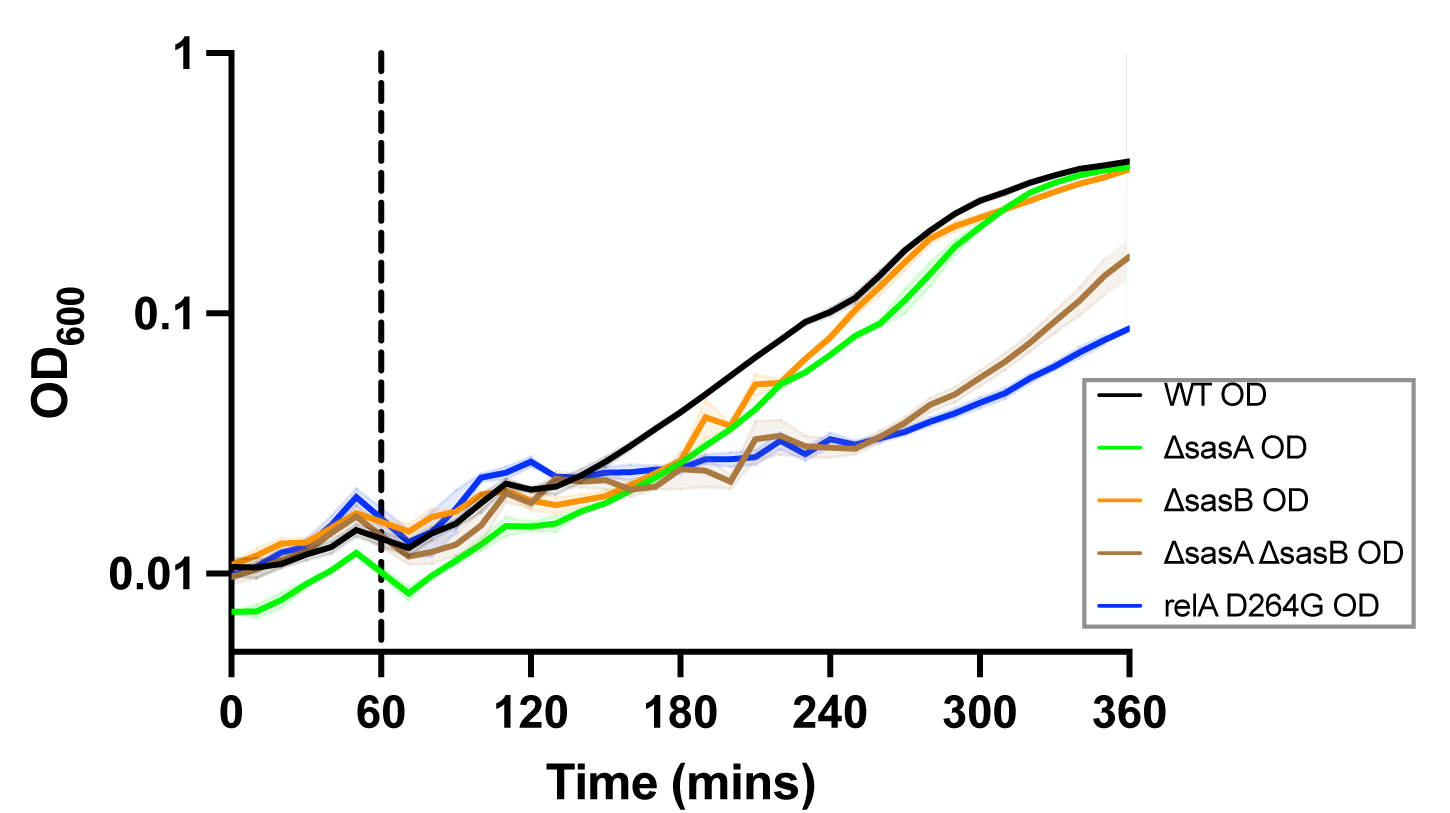
Synthases necessary for growth recovery from nutrient downshift. Luminescence (RLU/OD_600_) of RsFluc measured post nutrient downshift at T_60_ (dashed line) in WT (black, JDB4496), Δ*sasA* (green, JDB4515), Δ*sasB* (orange, JDB4516), Δ*sasA*Δ*sasB* (gold, JDB4508), and *relA-D264G* (red, JDB4741) backgrounds.

**Fig. S7.**
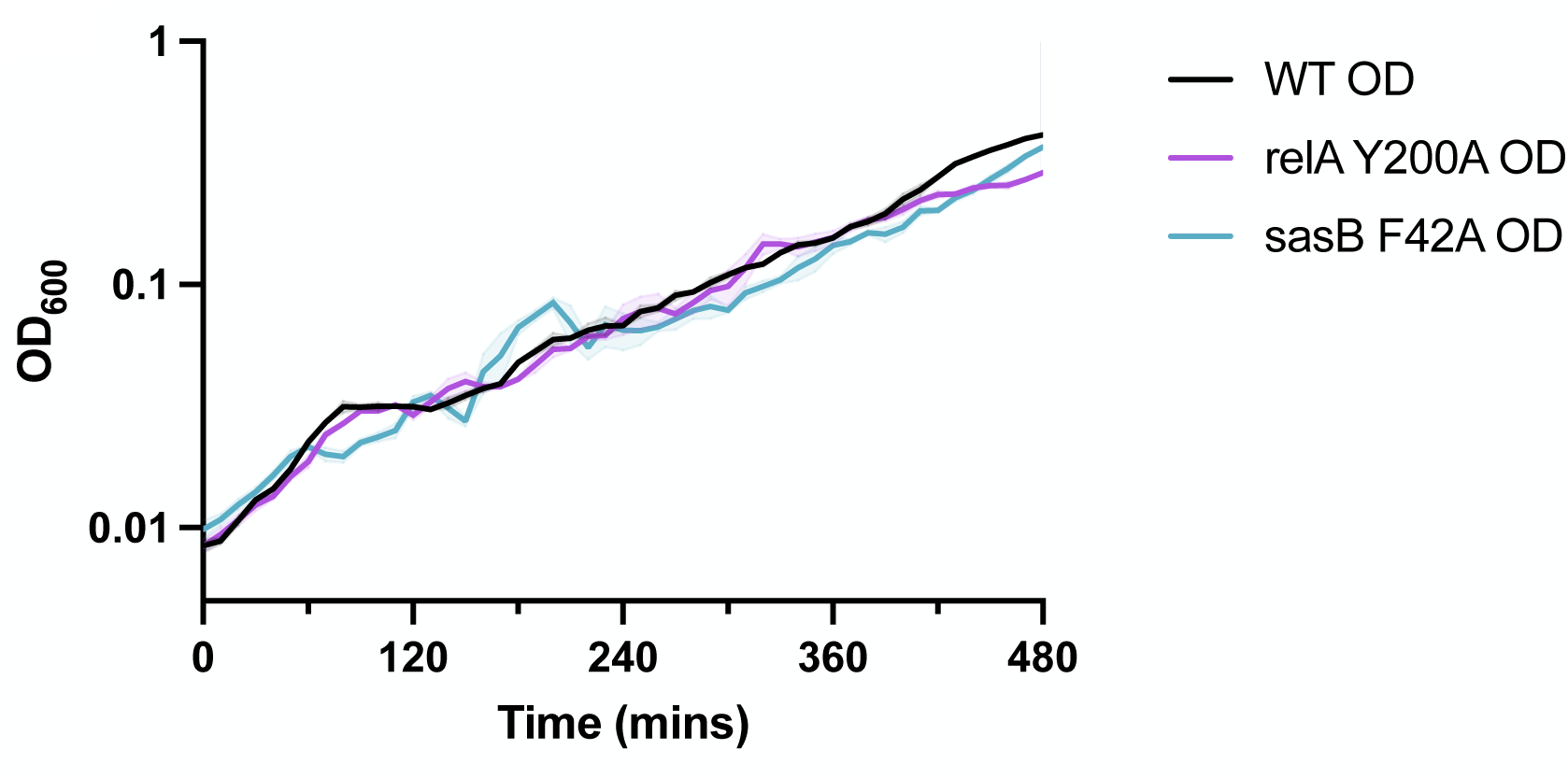
Growth of (p)ppGpp allosteric synthase mutants. Growth (OD_600_) of wildtype (black, JDB4496), *relA-Y200A* (fuschia, JDB4528), and *sasB-F42A* (sky blue, JDB4711) strains.

**Fig. S8.**
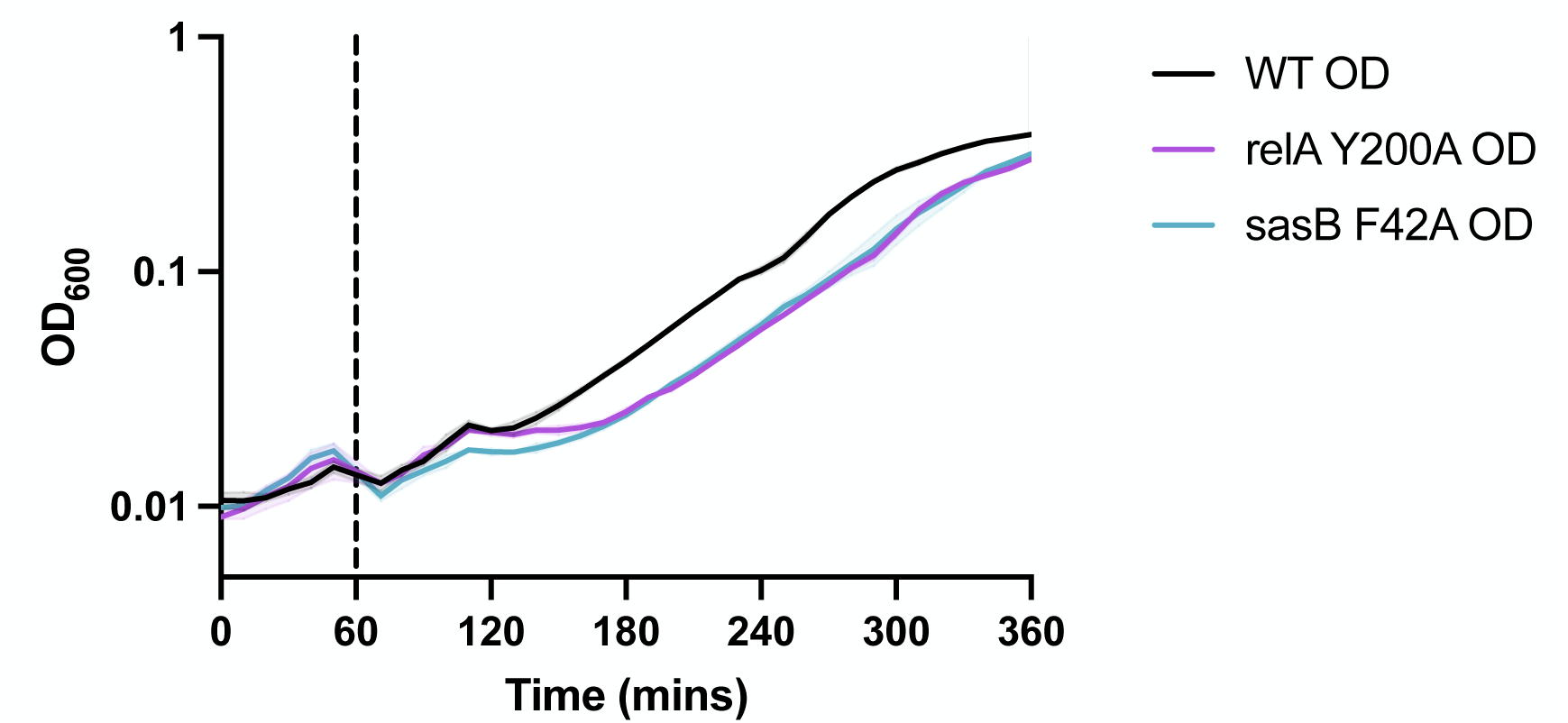
Allosteric regulation affects recovery post downshift. Growth (OD_600_) post nutrient downshift at T_60_, (dashed line) in wildtype (black, JDB4496), *relA-Y200A* (fuschia, JDB4528), and *sasB-F42A* (sky blue, JDB4711) strains.

**Fig. S9.**
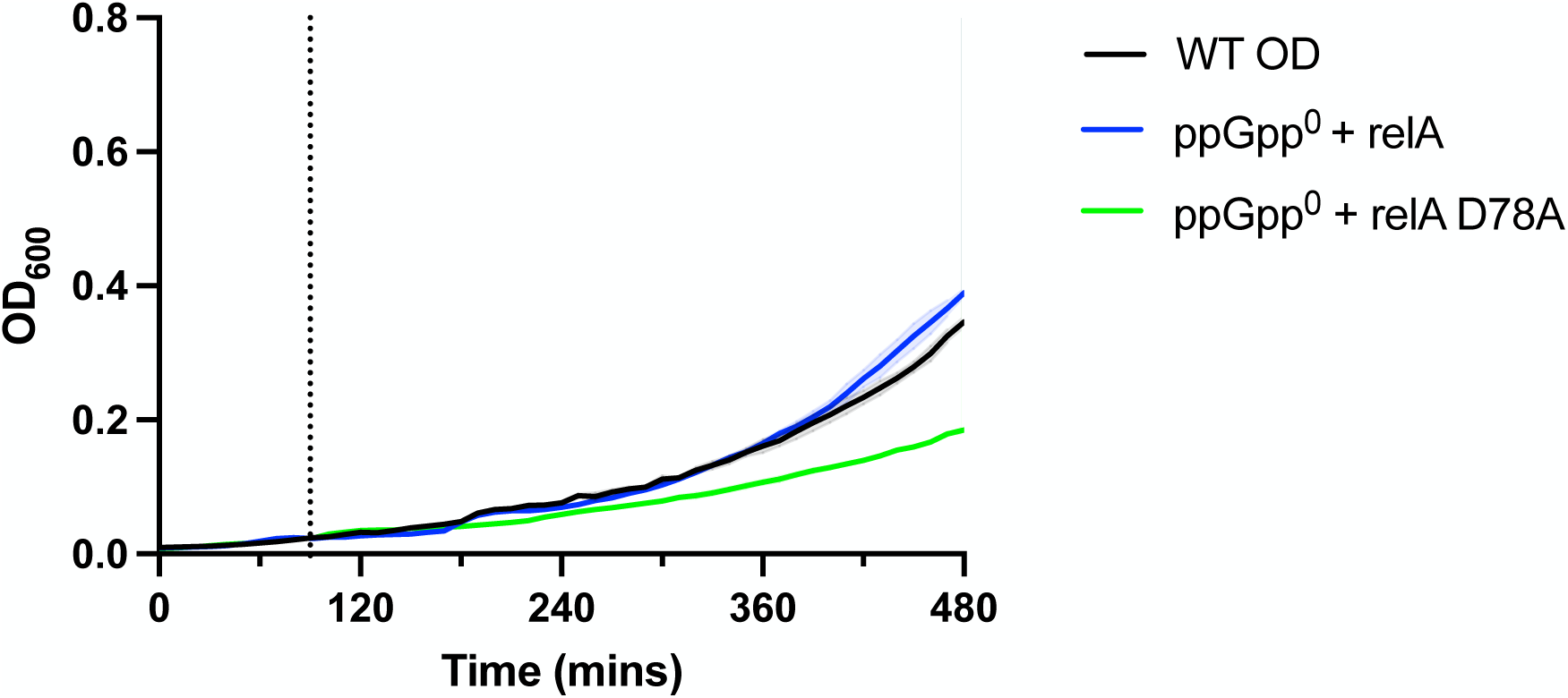
Growth of inducible WT and RelA-D78A. Growth (OD_600_) before and after 5 µg/mL bacitracin addition (dashed line) in WT (black, JDB4496), (p)ppGpp^0^ with inducible WT *relA* (blue, JDB4675), and (p)ppGpp^0^ with inducible *relA-D78A* (green, JDB4676) backgrounds.

**Fig. S10.**
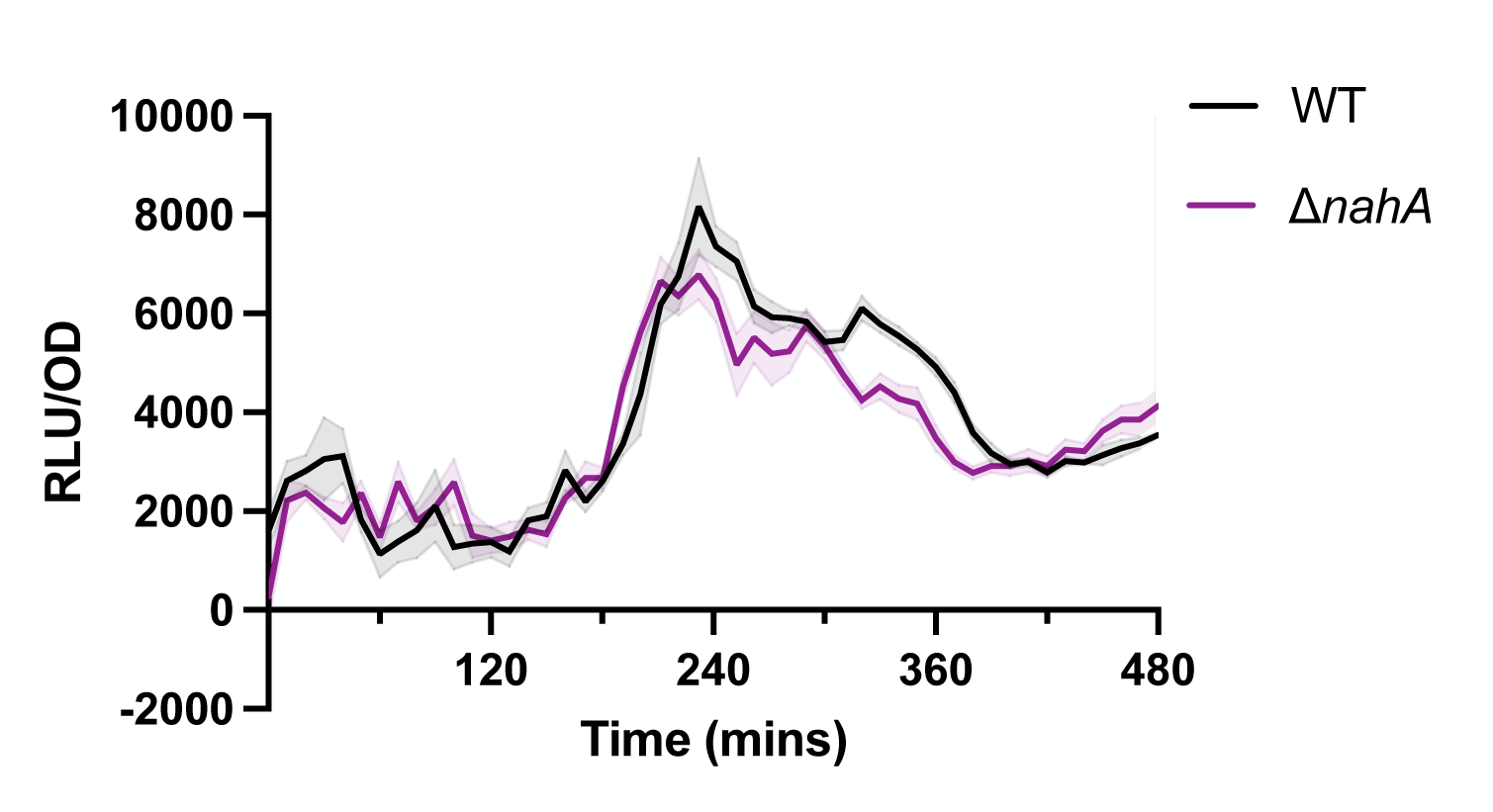
NahA contribution to RsFluc activity. Luminescence (RLU/OD_600_) of RsFluc in wildtype (black, JDB4496) and Δ*nahA* (purple, JDB4567) backgrounds.

**Fig. S11.**
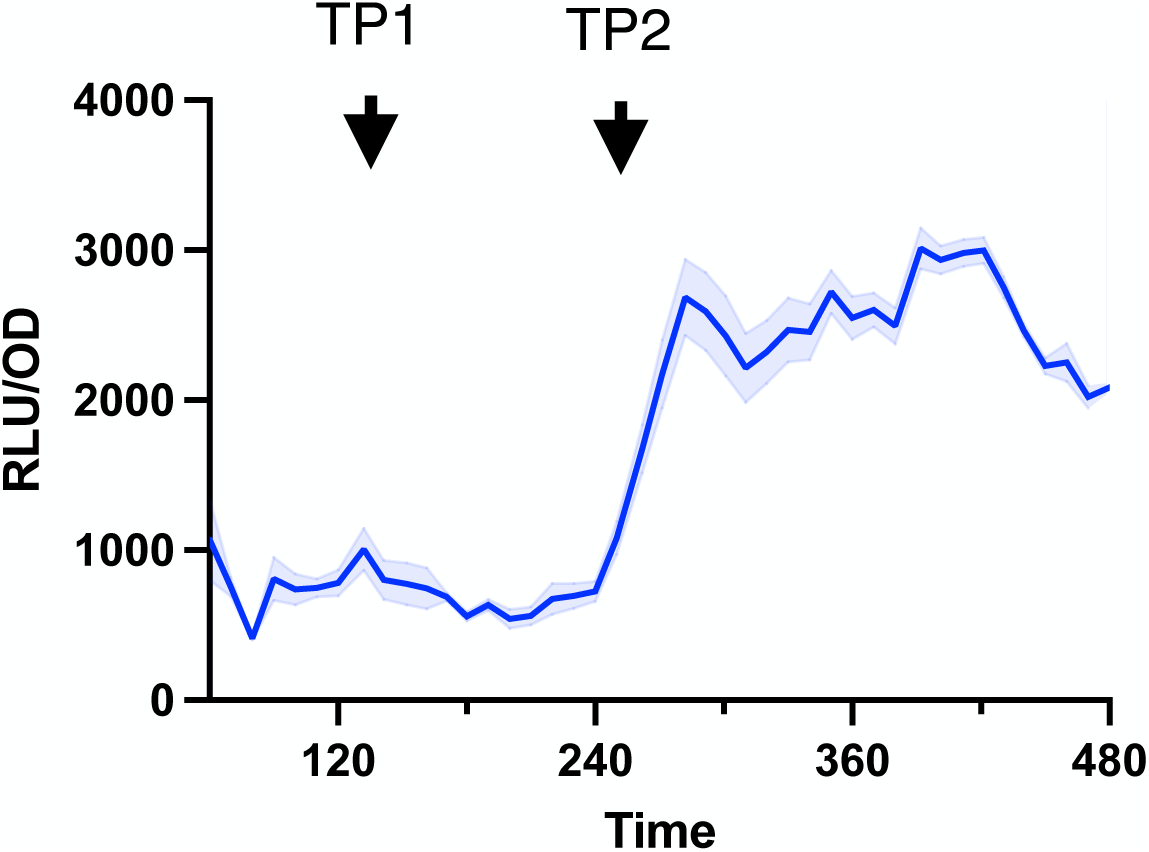
Time points for OPP labeling. Luminescence (RLU/OD_600_) of RsFluc in wildtype (blue, JDB4496). Samples were taken at time points TP1 (120 min) and TP2 (240 min).

**Table S1.** Strains.

| Strain | Genotype | Source |
| --- | --- | --- |
| JDB3 | *Prototroph (PY79)* | Lab collection |
| JDB1772 | *trpC2* | Lab collection |
| JDB4294 | *trpC2 relA::erm sasA(ywaC)::kan yjbM::tet* | (59) |
| JDB4300 | *trpC2 relA-Y308A* | (59) |
| JDB4440 | *trpC2 yjbM-F42A* | (59) |
| JDB4525 | *trpC2 relA-Y200A* | This study |
| JDB4729 | *trpC2 relA D264G* | This study |
| JDB4496 | *trpC2 sacA::Phyperspank-ilvE riboswitch-luciferase cmR* | This study |
| JDB4512 | *relA::erm ywaC::kan yjbM::tet sacA::Phyperspank-D. hafniense ilvE riboswitch-luciferase-cmR* | This study |
| JDB4631 | *trpC2 sacA::Phyperspank-M9+M11 riboswitch-luciferase-cmR* | This study |
| JDB4524 | *trpC2 sacA::Phyperspank-luciferase-cmR* | This study |
| JDB4599 | *trpC2 sacA::Phyperspank-M9 riboswitch-luciferase-cm* | This study |
| JDB4522 | *trpC2 sacA::Phyperspank-M11 riboswitch-firefly luciferase-cm* | This study |
| JDB4730 | *trpC2 sacA::Phyperspank-Clostridiales natA riboswitch-luciferase-cmR* | This study |
| JDB4731 | *trpC2 sacA::Phyperspank-Oxobacter pfennigii livK riboswitch-luciferase-cmR* | This study |
| JDB4508 | *trpC2 ywaC::kan yjbM::tet sacA::Phyperspank-D. hafniense ilvE riboswitch-luciferase-cmR* | This study |
| JDB4515 | *trpC2 ywaC::kan sacA::Phyperspank-D. hafniense ilvE riboswitch-firefly luciferase-cmR* | This study |
| JDB4516 | *trpC2 yjbM::tet sacA::Phyperspank-D. hafniense ilvE riboswitch-firefly luciferase-cmR* | This study |
| JDB4741 | *trpC2 relA D264G sacA::Phyperspank-D. hafniense ilvE riboswitch-luciferase-cmR* | This study |
| JDB4568 | *trpC2 ykuL::kan sacA::Phyperspank-D. hafniense ilvE riboswitch-luciferase-cmR* | This study |
| JDB4675 | *trpC2 relA::erm ywaC::kan yjbM::tet sacA::Phyperspank-D. hafniense ilvE riboswitch-luciferase-cmR amyE::Plial-relA-specR* | This study |
| JDB4676 | *trpC2 relA::erm ywaC::kan yjbM::tet sacA::Phyperspank-D. hafniense ilvE riboswitch-luciferase-cmR amyE::Plial-relA D78A-specR* | This study |
| JDB4567 | *trpC2 yvcl::erm sacA::Phyperspank-D. hafniense ilvE riboswitch-luciferase-cmR* | This study |
| JDB4711 | *trpC2 yjbM F42A sacA::Phyperspank-D. hafniense ilvE riboswitch-luciferase-cmR* | This study |
| JDB4528 | *trpC2 relA Y200A sacA::Phyperspank-D. hafniense ilvE riboswitch-luciferase-cmR* | This study |
| JDB4656 | *PY79 sacA::Phyperspank-D. hafniense ilvE riboswitch-luciferase-cmR* | This study |
| JDB4657 | *PY79 sacA::Phyperspank-M9+M11 riboswitch-luciferase-cmR* | This study |
| JDB4759 | *trpC2 sacA::PserA-luciferase-cmR* | This study |
| JDB4765 | *trpC2 sacA::PhomA-luciferase-cmR* | This study |
| JDB4792 | *trpC2 sacA::PilvB-luciferase-cmR* | This study |
| JDB4798 | *trpC2 sacA::PmetE-luciferase-cmR* | This study |
| JDB4793 | *trpC2 relA D264G sacA::PserA-luciferase-cmR* | This study |
| JDB4794 | *trpC2 relA D264G sacA::PhomA-luciferase-cmR* | This study |
| JDB4795 | *trpC2 relA D264G sacA::PilvB-luciferase-cmR* | This study |
| JDB4799 | *trpC2 relA D264G sacA::PmetE-luciferase-cmR* | This study |
| JDB4803 | *trpC2 sacA::PpurE-luciferase-cmR* | This study |
| JDB4804 | *trpC2 relA Y308A ywaC::kan yjbM::tet sacA::PpurE-firefly -cmR* | This study |

**Table S2.**
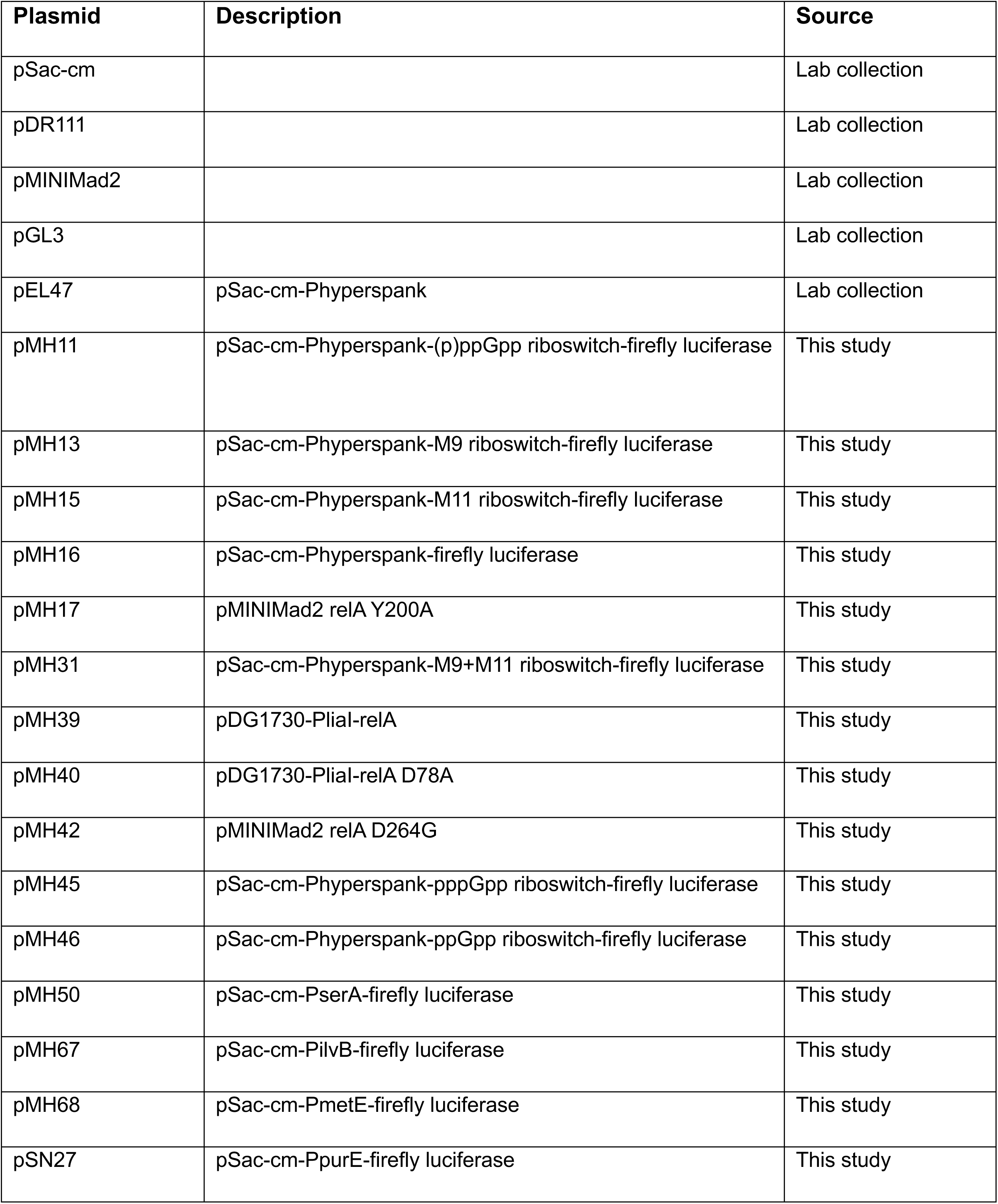
Plasmids.

**Table S3.**
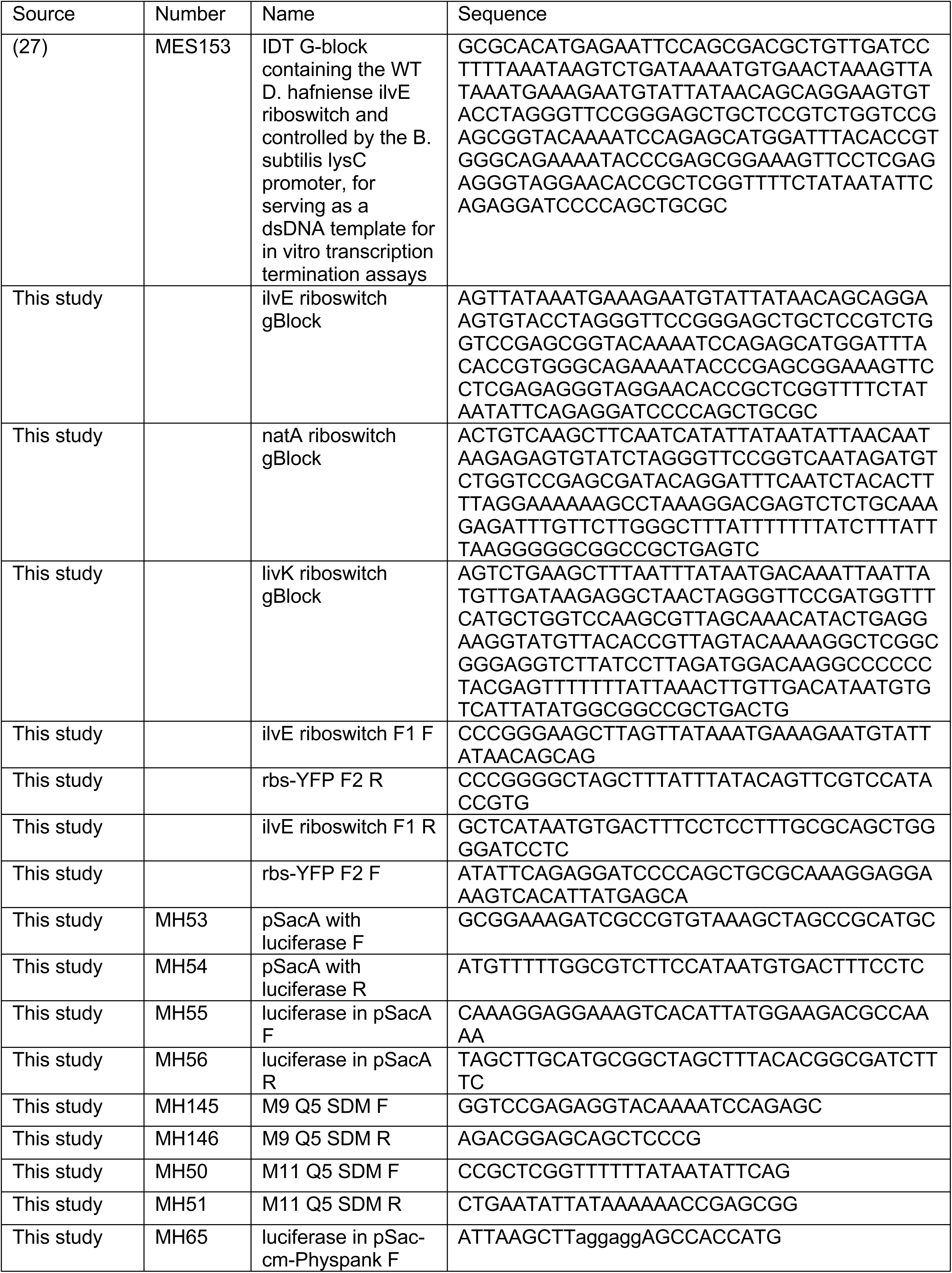
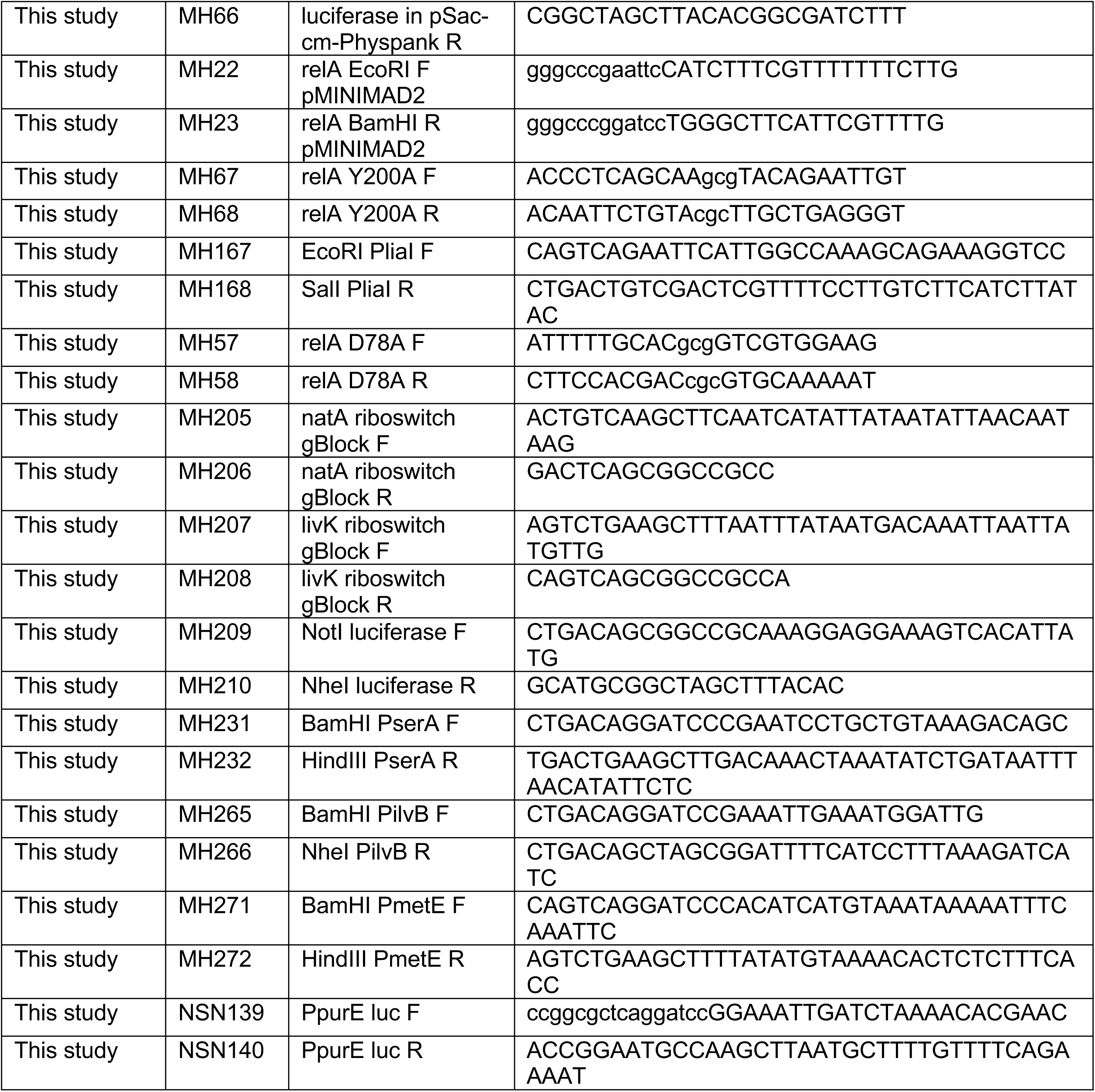
Oligos.

## References

1. Anderson BW, Fung DK, Wang JD. Regulatory Themes and Variations by the Stress-Signaling Nucleotide Alarmones (p)ppGpp in Bacteria. Annu Rev Genet. 2021;55:115–33.

2. Bange G, Brodersen DE, Liuzzi A, Steinchen W. Two P or Not Two P: Understanding Regulation by the Bacterial Second Messengers (p)ppGpp. Annu Rev Microbiol. 2021;75:383–406.

3. Schafer H, Beckert B, Frese CK, Steinchen W, Nuss AM, Beckstette M, et al. The alarmones (p)ppGpp are part of the heat shock response of Bacillus subtilis. PLoS Genet. 2020;16(3):e1008275.

4. Atkinson GC, Tenson T, Hauryliuk V. The RelA/SpoT homolog (RSH) superfamily: distribution and functional evolution of ppGpp synthetases and hydrolases across the tree of life. PLoS One. 2011;6(8):e23479.

5. Haseltine WA, Block R. Synthesis of guanosine tetra- and pentaphosphate requires the presence of a codon-specific, uncharged transfer ribonucleic acid in the acceptor site of ribosomes. Proc Natl Acad Sci U S A. 1973;70(5):1564–8.

6. Hogg T, Mechold U, Malke H, Cashel M, Hilgenfeld R. Conformational antagonism between opposing active sites in a bifunctional RelA/SpoT homolog modulates (p)ppGpp metabolism during the stringent response [corrected]. Cell. 2004;117(1):57–68.

7. Nanamiya H, Kasai K, Nozawa A, Yun C-S, Narisawa T, Murakami K, et al. Identification and functional analysis of novel (p)ppGpp synthetase genes in Bacillus subtilis. Mol Microbiol. 2008;67(2):291–304.

8. Brown A, Fernandez IS, Gordiyenko Y, Ramakrishnan V. Ribosome-dependent activation of stringent control. Nature. 2016;534(7606):277-80.

9. Loveland AB, Bah E, Madireddy R, Zhang Y, Brilot AF, Grigorieff N, et al. Ribosome*RelA structures reveal the mechanism of stringent response activation. Elife. 2016;5.

10. Takada H, Roghanian M, Caballero-Montes J, Van Nerom K, Jimmy S, Kudrin P, et al. Ribosome association primes the stringent factor Rel for tRNA-dependent locking in the A-site and activation of (p)ppGpp synthesis. Nucleic Acids Res. 2021;49(1):444–57.

11. Shyp V, Tankov S, Ermakov A, Kudrin P, English BP, Ehrenberg M, et al. Positive allosteric feedback regulation of the stringent response enzyme RelA by its product. EMBO Rep. 2012;13(9):835–9.

12. Steinchen W, Schuhmacher JS, Altegoer F, Fage CD, Srinivasan V, Linne U, et al. Catalytic mechanism and allosteric regulation of an oligomeric (p)ppGpp synthetase by an alarmone. Proc Natl Acad Sci U S A. 2015;112(43):13348–53.

13. Diez S, Hydorn M, Whalen A, Dworkin J. Crosstalk between guanosine nucleotides regulates cellular heterogeneity in protein synthesis during nutrient limitation. PLoS Genet. 2022;18(5):e1009957.

14. Fernandez-Coll L, Cashel M. Contributions of SpoT Hydrolase, SpoT Synthetase, and RelA Synthetase to Carbon Source Diauxic Growth Transitions in Escherichia coli. Front Microbiol. 2018;9:1802.

15. Lee JW, Park YH, Seok YJ. Rsd balances (p)ppGpp level by stimulating the hydrolase activity of SpoT during carbon source downshift in Escherichia coli. Proc Natl Acad Sci U S A. 2018;115(29):E6845–E54.

16. Raskin DM, Judson N, Mekalanos JJ. Regulation of the stringent response is the essential function of the conserved bacterial G protein CgtA in Vibrio cholerae. Proc Natl Acad Sci U S A. 2007;104(11):4636–41.

17. Fang M, Bauer CE. Regulation of stringent factor by branched-chain amino acids. Proc Natl Acad Sci U S A. 2018;115(25):6446–51.

18. Roghanian M, Van Nerom K, Takada H, Caballero-Montes J, Tamman H, Kudrin P, et al. (p)ppGpp controls stringent factors by exploiting antagonistic allosteric coupling between catalytic domains. Mol Cell. 2021;81(16):3310–22 e6.

19. Sinha AK, Winther KS. The RelA hydrolase domain acts as a molecular switch for (p)ppGpp synthesis. Commun Biol. 2021;4(1):434.

20. Pausch P, Abdelshahid M, Steinchen W, Schafer H, Gratani FL, Freibert SA, et al. Structural Basis for Regulation of the Opposing (p)ppGpp Synthetase and Hydrolase within the Stringent Response Orchestrator Rel. Cell Rep. 2020;32(11):108157.

21. Parker DJ, Lalanne JB, Kimura S, Johnson GE, Waldor MK, Li GW. Growth-Optimized Aminoacyl-tRNA Synthetase Levels Prevent Maximal tRNA Charging. Cell Syst. 2020;11(2):121–30 e6.

22. Fernandez-Coll L, Cashel M. Possible Roles for Basal Levels of (p)ppGpp: Growth Efficiency Vs. Surviving Stress. Front Microbiol. 2020;11:592718.

23. Varik V, Oliveira SRA, Hauryliuk V, Tenson T. HPLC-based quantification of bacterial housekeeping nucleotides and alarmone messengers ppGpp and pppGpp. Sci Rep. 2017;7(1):11022.

24. Patacq C, Chaudet N, Letisse F. Absolute Quantification of ppGpp and pppGpp by Double-Spike Isotope Dilution Ion Chromatography-High-Resolution Mass Spectrometry. Anal Chem. 2018;90(18):10715–23.

25. Zborníková E, Knejzlík Z, Hauryliuk V, Krásný L, Rejman D. Analysis of nucleotide pools in bacteria using HPLC-MS in HILIC mode. Talanta. 2019;205:120161.

26. Fung DK, Yang J, Stevenson DM, Amador-Noguez D, Wang JD. Small Alarmone Synthetase SasA Expression Leads to Concomitant Accumulation of pGpp, ppApp, and AppppA in Bacillus subtilis. Front Microbiol. 2020;11:2083.

27. Sherlock ME, Sudarsan N, Breaker RR. Riboswitches for the alarmone ppGpp expand the collection of RNA-based signaling systems. Proc Natl Acad Sci U S A. 2018;115(23):6052–7.

28. Mirouze N, Prepiak P, Dubnau D. Fluctuations in spo0A transcription control rare developmental transitions in Bacillus subtilis. PLoS Genet. 2011;7(4):e1002048.

29. Jagodnik J, Tjaden B, Ross W, Gourse RL. Identification and characterization of RNA binding sites for (p)ppGpp using RNA-DRaCALA. Nucleic Acids Res. 2023;51(2):852–69.

30. Wendrich TM, Marahiel MA. Cloning and characterization of a relA/spoT homologue from Bacillus subtilis. Mol Microbiol. 1997;26(1):65–79.

31. Sorensen MA. Charging levels of four tRNA species in Escherichia coli Rel(+) and Rel(-) strains during amino acid starvation: a simple model for the effect of ppGpp on translational accuracy. J Mol Biol. 2001;307(3):785–98.

32. Zhu M, Dai X. Stringent response ensures the timely adaptation of bacterial growth to nutrient downshift. Nat Commun. 2023;14(1):467.

33. Dittmar KA, Sorensen MA, Elf J, Ehrenberg M, Pan T. Selective charging of tRNA isoacceptors induced by amino-acid starvation. EMBO Rep. 2005;6(2):151–7.

34. Fung DK, Barra JT, Schroeder JW, Ying D, Wang JD. A shared alarmone-GTP switch underlies triggered and spontaneous persistence. bioRxiv. 2020:2020.03.22.002139.

35. Hughes J, Mellows G. Inhibition of isoleucyl-transfer ribonucleic acid synthetase in Escherichia coli by pseudomonic acid. Biochem J. 1978;176(1):305–18.

36. Cassels R, Oliva B, Knowles D. Occurrence of the regulatory nucleotides ppGpp and pppGpp following induction of the stringent response in staphylococci. J Bacteriol. 1995;177(17):5161–5.

37. Traxler MF, Chang DE, Conway T. Guanosine 3’,5’-bispyrophosphate coordinates global gene expression during glucose-lactose diauxie in Escherichia coli. Proc Natl Acad Sci U S A. 2006;103(7):2374–9.

38. Monod J. Recherches sur la Croissance des Cultures Bactériennes. Paris.: Hermann.; 1941.

39. Anderson BW, Schumacher MA, Yang J, Turdiev A, Turdiev H, Schroeder JW, et al. The nucleotide messenger (p)ppGpp is an anti-inducer of the purine synthesis transcription regulator PurR in Bacillus. Nucleic Acids Res. 2022;50(2):847–66.

40. Srivatsan A, Han Y, Peng J, Tehranchi AK, Gibbs R, Wang JD, et al. High-precision, whole-genome sequencing of laboratory strains facilitates genetic studies. PLoS Genet. 2008;4(8):e1000139.

41. Radeck J, Kraft K, Bartels J, Cikovic T, Durr F, Emenegger J, et al. The Bacillus BioBrick Box: generation and evaluation of essential genetic building blocks for standardized work with Bacillus subtilis. J Biol Eng. 2013;7(1):29.

42. Yang J, Anderson BW, Turdiev A, Turdiev H, Stevenson DM, Amador-Noguez D, et al. The nucleotide pGpp acts as a third alarmone in Bacillus, with functions distinct from those of (p) ppGpp. Nat Commun. 2020;11(1):5388.

43. Kriel A, Brinsmade SR, Tse JL, Tehranchi AK, Bittner AN, Sonenshein AL, et al. GTP dysregulation in Bacillus subtilis cells lacking (p)ppGpp results in phenotypic amino acid auxotrophy and failure to adapt to nutrient downshift and regulate biosynthesis genes. J Bacteriol. 2014;196(1):189–201.

44. Diez S, Ryu J, Caban K, Gonzalez RL, Jr., Dworkin J. The alarmones (p)ppGpp directly regulate translation initiation during entry into quiescence. Proc Natl Acad Sci U S A. 2020;117(27):15565–72.

45. Scott M, Gunderson CW, Mateescu EM, Zhang Z, Hwa T. Interdependence of cell growth and gene expression: origins and consequences. Science. 2010;330(6007):1099-102.

46. Gaca AO, Kajfasz JK, Miller JH, Liu K, Wang JD, Abranches J, et al. Basal levels of (p)ppGpp in Enterococcus faecalis: the magic beyond the stringent response. mBio. 2013;4(5):e00646–13.

47. Imholz NCE, Noga MJ, van den Broek NJF, Bokinsky G. Calibrating the Bacterial Growth Rate Speedometer: A Re-evaluation of the Relationship Between Basal ppGpp, Growth, and RNA Synthesis in Escherichia coli. Front Microbiol. 2020;11:574872.

48. Ababneh QO, Herman JK. RelA inhibits Bacillus subtilis motility and chaining. J Bacteriol. 2015;197(1):128–37.

49. Trinquier A, Ulmer JE, Gilet L, Figaro S, Hammann P, Kuhn L, et al. tRNA Maturation Defects Lead to Inhibition of rRNA Processing via Synthesis of pppGpp. Mol Cell. 2019;74(6):1227–38 e3.

50. Gallant J, Irr J, Cashel M. The mechanism of amino acid control of guanylate and adenylate biosynthesis. J Biol Chem. 1971;246(18):5812–6.

51. Lopez JM, Dromerick A, Freese E. Response of guanosine 5’-triphosphate concentration to nutritional changes and its significance for Bacillus subtilis sporulation. J Bacteriol. 1981;146(2):605–13.

52. Kriel A, Bittner AN, Kim SH, Liu K, Tehranchi AK, Zou WY, et al. Direct regulation of GTP homeostasis by (p)ppGpp: a critical component of viability and stress resistance. Mol Cell. 2012;48(2):231–41.

53. Tagami K, Nanamiya H, Kazo Y, Maehashi M, Suzuki S, Namba E, et al. Expression of a small (p)ppGpp synthetase, YwaC, in the (p)ppGpp(0) mutant of Bacillus subtilis triggers YvyD-dependent dimerization of ribosome. Microbiologyopen. 2012;1(2):115–34.

54. Patacq C, Chaudet N, Letisse F. Crucial Role of ppGpp in the Resilience of Escherichia coli to Growth Disruption. mSphere. 2020;5(6).

55. Harshman RB, Yamazaki H. Formation of ppGpp in a relaxed and stringent strain of Escherichia coli during diauxie lag. Biochemistry. 1971;10(21):3980–2.

56. Xiao H, Kalman M, Ikehara K, Zemel S, Glaser G, Cashel M. Residual guanosine 3’,5’-bispyrophosphate synthetic activity of relA null mutants can be eliminated by spoT null mutations. J Biol Chem. 1991;266(9):5980–90.

57. Solopova A, van Gestel J, Weissing FJ, Bachmann H, Teusink B, Kok J, et al. Bet-hedging during bacterial diauxic shift. Proc Natl Acad Sci U S A. 2014;111(20):7427–32.

58. Steinchen W, Vogt MS, Altegoer F, Giammarinaro PI, Horvatek P, Wolz C, et al. Structural and mechanistic divergence of the small (p)ppGpp synthetases RelP and RelQ. Sci Rep. 2018;8(1):2195.

59. Diez S, Ryu J, Caban K, Gonzalez RL, Dworkin J. The alarmones (p)ppGpp directly regulate translation initiation during entry into quiescence. Proc Natl Acad Sci U S A. 2020;117(27):15565–72.

60. Dworkin J. Understanding the Stringent Response: Experimental Context Matters. mBio. 2023;14(1):e0340422.

61. Gourse RL, Chen AY, Gopalkrishnan S, Sanchez-Vazquez P, Myers A, Ross W. Transcriptional Responses to ppGpp and DksA. Annu Rev Microbiol. 2018;72:163–84.

62. Weiss CA, Hoberg JA, Liu K, Tu BP, Winkler WC. Single-Cell Microscopy Reveals That Levels of Cyclic di-GMP Vary among Bacillus subtilis Subpopulations. J Bacteriol. 2019;201(16).

63. Kviatkovski I, Zhong Q, Vaidya S, Grundling A. Identification of novel genetic factors that regulate c-di-AMP production in Staphylococcus aureus using a riboswitch-based biosensor. mSphere. 2024:e0032124.

64. Sun Z, Wu R, Zhao B, Zeinert R, Chien P, You M. Live-Cell Imaging of Guanosine Tetra- and Pentaphosphate (p)ppGpp with RNA-based Fluorescent Sensors*. Angew Chem Int Ed Engl. 2021;60(45):24070–4.

65. Gaynor EC, Wells DH, MacKichan JK, Falkow S. The Campylobacter jejuni stringent response controls specific stress survival and virulence-associated phenotypes. Mol Microbiol. 2005;56(1):8–27.

66. Prusa J, Zhu DX, Stallings CL. The stringent response and Mycobacterium tuberculosis pathogenesis. Pathog Dis. 2018;76(5).

67. Dutta NK, Klinkenberg LG, Vazquez MJ, Segura-Carro D, Colmenarejo G, Ramon F, et al. Inhibiting the stringent response blocks Mycobacterium tuberculosis entry into quiescence and reduces persistence. Sci Adv. 2019;5(3):eaav2104.

68. Schofield WB, Zimmermann-Kogadeeva M, Zimmermann M, Barry NA, Goodman AL. The Stringent Response Determines the Ability of a Commensal Bacterium to Survive Starvation and to Persist in the Gut. Cell Host Microbe. 2018;24(1):120–32 e6.

69. Ontai-Brenning A, Hamchand R, Crawford JM, Goodman AL. Gut microbes modulate (p)ppGpp during a time-restricted feeding regimen. mBio. 2023;14(6):e0190723.

70. Foucault ML, Thomas L, Goussard S, Branchini BR, Grillot-Courvalin C. In vivo bioluminescence imaging for the study of intestinal colonization by Escherichia coli in mice. Appl Environ Microbiol. 2010;76(1):264–74.

71. Spira B, Ospino K. Diversity in E. coli (p)ppGpp Levels and Its Consequences. Front Microbiol. 2020;11:1759.

72. McCann JR, Rawls JF. Essential Amino Acid Metabolites as Chemical Mediators of Host-Microbe Interaction in the Gut. Annu Rev Microbiol. 2023;77:479–97.

73. Mee MT, Collins JJ, Church GM, Wang HH. Syntrophic exchange in synthetic microbial communities. Proc Natl Acad Sci U S A. 2014;111(20):E2149–56.

74. McKinlay JB. Are Bacteria Leaky? Mechanisms of Metabolite Externalization in Bacterial Cross-Feeding. Annu Rev Microbiol. 2023;77:277–97.

75. Pherribo GJ, Taga ME. Bacteriophage-mediated lysis supports robust growth of amino acid auxotrophs. ISME J. 2023;17(10):1785–8.

76. Middleton R, Hofmeister A. New shuttle vectors for ectopic insertion of genes into Bacillus subtilis. Plasmid. 2004;51(3):238–45.

77. Popp PF, Dotzler M, Radeck J, Bartels J, Mascher T. The Bacillus BioBrick Box 2.0: expanding the genetic toolbox for the standardized work with Bacillus subtilis. Sci Rep. 2017;7(1):15058.

78. Patrick JE, Kearns DB. Laboratory strains of Bacillus subtilis do not exhibit swarming motility. J Bacteriol. 2009;191(22):7129–33.

79. Ducret A, Quardokus EM, Brun YV. MicrobeJ, a tool for high throughput bacterial cell detection and quantitative analysis. Nat Microbiol. 2016;1(7):16077.

